# Controlling load-dependent contractility of the heart at the single molecule level

**DOI:** 10.1101/258020

**Authors:** Chao Liu, Masataka Kawana, Dan Song, Kathleen M. Ruppel, James A. Spudich

## Abstract

Concepts in molecular tension sensing in biology are growing and have their origins in studies of muscle contraction. In the heart muscle, a key parameter of contractility is the detachment rate from actin of myosin, which determines the time that myosin is bound to actin in a force-producing state and, importantly, depends on the load (force) against which myosin works. Here, we measure the detachment rate of single molecules of human β-cardiac myosin and its load dependence. We find that both can be modulated by both small molecule compounds and cardiomyopathy-causing mutations. Furthermore, effects of mutations can be reversed by introducing appropriate compounds. Our results suggest that activating vs. inhibitory perturbations of cardiac myosin are discriminated by the aggregate result on duty ratio, average force, and ultimately average power output and that cardiac contractility can be controlled by tuning the load-dependent kinetics of single myosin molecules.

## Main

The ability to sense and respond to force is a fundamental feature observed across all scales of biology, from tissues^1,2^, to cells^3,4^, and to individual molecules^5-8^. Muscle contraction is a classic example in which tension sensing of the tissue ^9^ has its roots in the individual molecular motors working in every muscle cell - the myosin molecule.

Detachment rate is a key kinetic parameter describing myosin’s actin-activated ATPase cycle because it determines the time myosin is bound to actin in a force-producing state. Importantly, this rate depends on the force (load) against which myosin works as demonstrated in single molecule optical trap experiments with cardiac ^10,11^, skeletal ^12^, smooth ^13^, and non-muscle myosins ^14-16^. The load-dependent detachment rate (*k*_*det*_(*F*)) is a fundamental characteristic of every individual myosin molecule. As such, it determines whether a non-muscle myosin is a force sensor (myosin Ib) or a transporter (myosins V, Ic), while for muscle myosins, it defines the identity of the particular muscle (skeletal, cardiac, smooth). Even within a particular muscle type, different isoforms have different load-dependent kinetics adapted to their specific function^17-19^. For instance, the beta isoform of cardiac muscle myosin is slower but has higher load-bearing ability than the alpha isoform ^20^, consistent with its dominant presence in the ventricles rather than the atria of the heart ^21^. The cardiac myosin identity is so specifically tuned such that the relative composition of the alpha and beta isoforms changes in the failing heart ^21^. Furthermore, replacement of one by the other had a detrimental effect in transgenic mice subjected to cardiovascular stress ^22^

Thus, it is apparent that significant differences in the amino acid sequences of different myosin types dramatically change *kdet*(*F*) to accommodate their molecular functions. However, it is not known whether small perturbations such as disease-causing mutations and small molecule drugs can control and alter the *k*_*det*_(*F*) in any specific myosin. Here, we address this question by measuring the effects of various small perturbations on the load-dependent kinetics of human β-cardiac myosin. Measurement of *k*_*det*_(*F*) of single striated muscle myosin molecules under physiological (~2 mM) ATP conditions is challenging due to their short binding lifetime.

Existing techniques, while able to measure *k*_*det*_(*F*), require fast feedback and/or are not efficient at obtaining a large number of events. In contrast, Harmonic Force Spectroscopy (HFS) presented a simple and efficient solution without the need for fast feedback ^11^. In the present study, we first demonstrate the robustness of the HFS method by measuring *k*_*det*_(*F*) for a population (N=36) of single molecules of truncated (residues 1-808) human β-cardiac subfragment-1 (short S1 or sS1), the motor domain of the myosin responsible for force production in the ventricles of the heart (hereafter referred to as myosin). The efficiency of this method allowed us to tackle the aforementioned biological question: how is myosin’s load-dependent kinetics controlled and affected by various biologically-relevant perturbations? We tested the extent to which small molecule compounds affect *k*_*det*_(*F*). These compounds include both allosteric activators and an inhibitor of cardiac myosin, including the potential heart failure drug omecamtiv mecarbil (OM) ^23^, and the potential active-site therapeutic 2-deoxy-ATP (dATP) ^24,25^. Next, we introduced mutations into β-cardiac sS1 that cause hypertrophic (HCM: D239N and H251N) or dilated (DCM: A223T, R237W and S532P) cardiomyopathies and measured their effects on *k*_*det*_(*F*). We show *that* the effects of mutations can be reversed by adding the appropriate small molecule compounds. In addition, we further investigated the mechanism of one of the compounds, OM, by measuring its dose-dependent effect on the detachment kinetics of myosin. Finally, our findings provide a new single molecule-level understanding of cardiac myosin’s power production under these various perturbations.

### Harmonic Force Spectroscopy enables simple and efficient measurement of load-dependent kinetics on single molecules of human β-cardiac myosin

In Harmonic Force Spectroscopy (HFS) optical trap experiments, the lifetimes of binding events between a single myosin motor and actin under various loads can be directly measured in saturating (2 mM) ATP conditions ^11^. The sample stage oscillates so that when myosin binds to actin, it experiences a sinusoidally varying load with a certain mean value (Fig. 1a). Unlike other methods which require feedback to apply a load ^10,12,13,26^, HFS does not. Oscillation serves two essential purposes: 1) A range of mean loads, both positive and negative, is automatically applied over the course of many events by virtue of the randomness of where myosin initially attaches to actin (Fig. 1a). 2) Oscillation allows for the detection of short binding events of cardiac myosin to actin, which were previously more difficult to detect than longer events. In the unattached state, the dumbbell oscillates π/2 ahead of the stage due to fluid motion, and in the attached state the dumbbell oscillates in phase and with a larger amplitude due to stronger association with the oscillating stage (Fig. 1b). Therefore, events are detected automatically based on a simultaneous increase in oscillation amplitude above a threshold and a decrease in phase below a threshold (Fig. 1b). In this way, HFS enables detection of short binding events (>5 ms, one period of stage oscillation) under load regardless of the length of an event.

**Figure 1:**
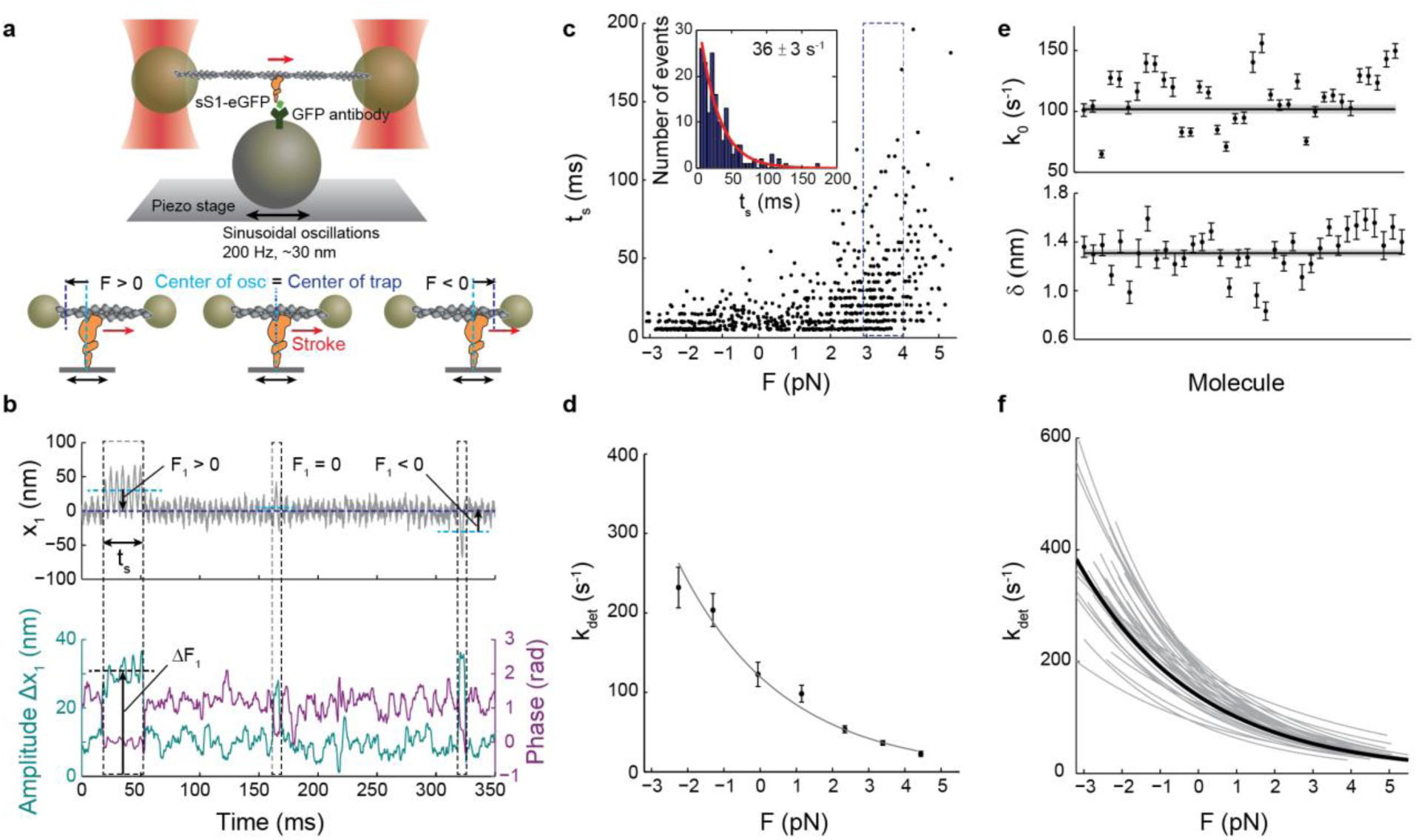
Load-dependent kinetics of single molecules of human β-cardiac myosin measured by harmonic force spectroscopy. **a**, Oscillations of the piezo stage on which myosin is attached in the three-bead optical trap system (top) apply a sinusoidal load (force) to myosin upon attachment to actin (bottom). **b**, Example trace of the position of one trapped bead, *x*_*1*_ (top). An increase in the amplitude of oscillation and a simultaneous decrease in the phase between the trapped bead and piezo stage (bottom) are the two criteria used in the automated detection of an event. Over the course of many individual binding events, a range of forces with mean *F*_*1*_(calculated as the position *xi* multiplied by the trap stiffness) is automatically applied as myosin attaches randomly along the actin dumbbell to experience both positive (resistive) or negative (assistive) loads ^11^. Three events are outlined, the first with positive *F*_*1*_, the second with *F*_*1*_ close to zero, and the third with negative *F*_*1*_ Each event experiences the oscillatory force with mean *Fi* and amplitude *ΔF*_*1*_, calculated as the amplitude *Δx*_*1*_ multiplied by the trap stiffness. Force *F* in Eqn. 1 is calculated as the sum of *F*_*1*_ and *F*_*2*_ from the second trapped bead, and the force amplitude *ΔF* is calculated as the sum of *ΔF*_*1*_ and *ΔF*_*2*_ (~3-4 pN). **c**, All events (N=729) for one example molecule. Duration of event (*t*_*s*_) is plotted against force. Events binned by force (dashed blue outline) have exponentially-distributed binding times from which the detachment rate at that force is determined by maximum likelihood estimation (MLE) (inset). The error on the rate is calculated from the variance of the MLE as the inverse Fisher information. **d**, The force-dependent Arrhenius equation with harmonic force correction (Eqn. 1) is fitted to the detachment rates at each force to yield the load-dependent kinetics curve for one molecule. The two fitting parameters are *k*_0_ = 84 ± 4 s^−1^, the rate at zero load, and *Δ* = 1.31 ± 0.03 nm, the distance to the transition state, which is a measure of force sensitivity. Errors on *k*_*0*_ and *Δ* are from the covariance matrix (inverse Fisher information matrix) for the parameters. **e**, *k*_*0*_ (top) and *Δ*(bottom) for all molecules measured (N=36). Their weighted means, *k*_*0*_ = 102 ± 4 s^−1^, *Δ* = 1.31 ± 0.03 nm, are shown as horizontal lines with s.e.m. in gray. **f**, Load-dependent kinetics curves for all molecules measured (gray), each described by a (*k*_0_, *δ*) pair in (e). Individual data points as in (d) are not shown for clarity. The curve corresponding to the weighted means of *k*_*0*_ and *δ* is shown in black. All experiments were done at saturating (2 mM) ATP.

In the analysis of time trace data for a single myosin molecule, binding events are first detected, each defined by the duration *t*_*s*_ and mean force *F* (Fig. 1c). Events are then binned in force increments of ~1 pN so that the detachment rate for each bin *k*_*det*_(*F*) can be calculated by a maximum likelihood estimation (MLE) on the exponentially-distributed bound times (Fig. 1c inset). Since the load experienced by myosin during an event is sinusoidal with amplitude *ΔF* and mean *F*, the dependence of the detachment rate *k*_*det*_ on force is given by the force-dependent Arrhenius equation with a harmonic force correction:

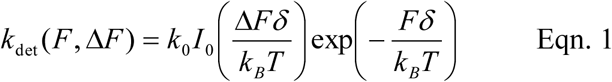

where *k*_*0*_ is the rate at zero load, δ is the distance to the transition state of the rate limiting step in the bound state (a measure of sensitivity to force), *k*_*B*_ is the Boltzmann constant, *T* is temperature, and *I*_*0*_ is the zeroth-order modified Bessel function of the first kind (to correct for the harmonic force). This equation is fitted to the detachment rates *k*_*det*_(*F*) with the parameters *k*_*0*_ and δ to obtain the loaded curve for a single myosin molecule (Fig. 1d).

Measurements of a population (N=36) of single molecules of human β-cardiac myosin yielded the mean values *k*_*0*_ = 102 ± 4 s^−1^ and δ = 1.31 ± 0.03 nm (standard error of mean, s.e.m.) (Fig. 1e-f), consistent with previously reported values ^10,11^. The HFS method allowed us to produce a load-dependent curve for each molecule rather than one from events aggregated across multiple molecules, therefore producing the distribution of the population and revealing inherent molecule-to-molecule differences (Fig. 1e-f).

### Allosteric effectors of human β-cardiac myosin slow down ensemble actin-sliding velocity

We investigated four allosteric activators (A2876 (1), C2981 (2), D3390 (3), OM), one allosteric inhibitor (F3345 (4)), and the active-site substrate dATP as small molecule effectors of cardiac myosin (Fig. 2a). dATP, a potential heart failure therapeutic ^24^, acts in place of ATP and has been shown to enhance cardiac myosin’s ATPase activity ^25^. A2876 (1), C2981 (2), and D3390 (3) were derived from hits discovered from an ATPase screen using bovine cardiac subfragment-1 (S1) and regulated thin filaments (purified bovine actin, tropomyosin, and troponins C, I, and T), and F3345 (4) was derived from a hit discovered from an ATPase screen using bovine myofibrils, all by MyoKardia, Inc. OM, an investigational drug in phase III clinical trials for heart failure ^23,27^ (GALACTIC-HF, www.clinicaltrials.gov NCT02929329), has been proposed to increase ensemble force by increasing myosin’s duty ratio, the fraction of time spent in the strongly-bound, force producing state ^20,28-30^. The rate of ADP release normally determines the strongly-bound state time, but surprisingly, OM was found to delay the power stroke ^30^ without affecting this rate ^23,28-30^. To clarify OM’s mechanism of action, we used HFS to directly measure whether the drug causes a single cardiac myosin molecule to stay bound longer to actin.

**Figure 2:**
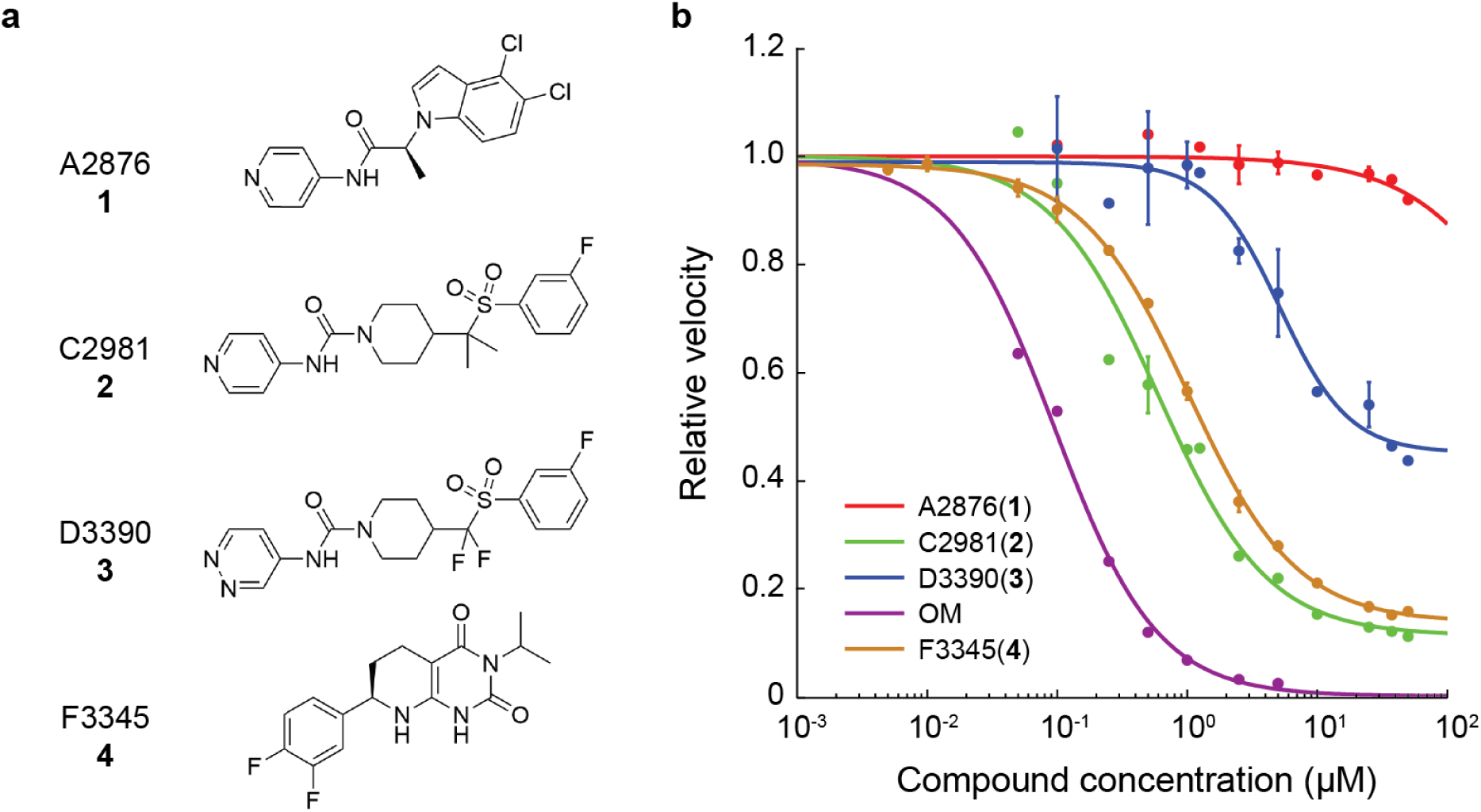
Dose-dependent effects of small molecule compounds on the actin-sliding velocity of human β-cardiac myosin in unloaded in vitro motility. **a**, Structures of the compounds used in this study. The structure of omecamtiv mecarbil (OM) is reported in Malik et al. ^23^. **b**, Dose-dependent effects of the compounds on the actin-sliding velocity of human β-cardiac myosin. Velocities are normalized to the value at zero compound concentration. Note that some data points have error bars smaller than the displayed point. Error bars represent s.e.m. between experiments, each tracking ~400 to ~1900 actin filament movements. Curves are 4-parameter logistic fits.

To determine the effects of the compounds when a single myosin molecule has a compound bound, a saturating concentration of compound was required. To determine this concentration, we measured the dose-dependent effects of the compounds on the actin-sliding velocities of human β-cardiac myosin in an in vitro motility assay ^20,31^. Surprisingly, both activators and inhibitors slowed down velocities, but to different extents (Fig. 2b). In particular, OM had the most dramatic effect with a median effective concentration (EC50) ~0.1 μM, in close agreement with previous measurements ^20,28,29^ Activator A2876 (1) had no effect on the velocity beyond that from DMSO alone (see methods).

Guided by the motility measurements, we performed subsequent single-molecule experiments that involve allosteric effectors at a concentration of 25 µM in 2% DMSO.

### Small molecule compounds modulate the load-dependent kinetics of single molecules of human β-cardiac myosin

We measured the effects of the small molecule compounds on the load-dependent detachment rates of single molecules of human β-cardiac myosin using HFS (Fig. 3). For clarity, we present only the average load-dependent kinetics curve for each condition in Fig. 3b (for curves of the individual molecules, see Extended Data Figs. 2 and 3). Values of *k*_*0*_ and *δ* are summarized in Table 1 and Extended Data Table 1. When dATP was used as myosin’s substrate, the rate *k*_*0*_increased (dATP: *k*_*0*_ = 168 ± 7 s^−1^ vs ATP: 102 ± 4 s^−1^) while *δ,* a measure of the sensitivity to force, did not change significantly (dATP: *δ* = 1.5 ± 0.1 nm vs ATP: *δ* = 1.31± 0.03 nm, p = 0.13), in close agreement with experiments performed using bovine cardiac heavy meromyosin (HMM) (Tomasic I., Liu C., Rodriguez H., Spudich J.A., Bartholomew Ingle S.R., manuscript in preparation). Compound A2876 (1) had no effect beyond that from DMSO. This result is consistent with observations that A2876 (1) has no effect on the actin-activated ATPase of purified bovine cardiac myosin and actin, but only activates myosin’s ATPase activity in the presence of regulated thin filaments, suggesting possible interactions with the tropomyosin-troponin complex (R. Anderson, MyoKardia unpublished results). The activators C2981 (2), D3390 (3), and OM decreased *k*_*0*_ dramatically (C2981 (2): *k*_*0*_ = 23 ± 1 s^−1^, D3390 (3): *k*_*0*_ = 55 ± 3 s^−1^, OM: *k*_*0*_ = 12 ± 1 s^−1^) while also decreasing *δ* (C2981 (2): *δ* = 0.79 ± 0.05 nm, D3390 (3): *δ* = 0.93 ± 0.03 nm, OM: *δ* =0.26 ± 0.05 nm) (Fig. 3a), resulting in a much lower and flatter curve that is less sensitive to force (Fig. 3b). All three of these activators probably interact directly with myosin, and lowering of both *k*_*0*_ and *δ* may be a general characteristic of myosin-associated activators. In contrast, the inhibitor F3345 (4) decreased the detachment rate (*k*_*0*_ = 36 ± 2 s^−1^) without changing the force sensitivity (*δ*= 1.27 ± 0.06 nm), suggesting a possible mechanism of action distinct from the activators. Taken together, we found that small molecule effectors can modulate the detachment rate and load sensitivity of cardiac myosin to various degrees.

**Table 1.** Summary of single molecule detachment kinetics (*k*_*0*_, *δ*) and actin-activated ATPase (*k*_*cat*_) data for human β-cardiac myosin with effects of small molecule compounds and cardiomyopathy-causing mutations. Values are mean ± s.e.m. The attachment rate *k*_*attach*_ is calculated by Eqn. 3, with propagated error. Values given in this table are used in Fig. 7. *k*_*cat*_ of mutants are obtained from other studies ^20,32^ (Ujfalusi Z., Vera C., Mijailovich S., Svicevic M., Choe Yu E., Kawana M., Ruppel K., Spudich J., Geeves M., and Leinwand L., *JBC* manuscript under review). In the case of dATP, 2 mM dATP was used in place of ATP. All other conditions used 2 mM ATP, 2% DMSO, and 25 μM compound concentration. 2-tailed unequal variances (for *k*_*0*_ and *δ*) and paired (for *k*_*cat*_) (see Methods) t-test performed on dATP vs ATP, DMSO vs WT, compounds + DMSO vs DMSO alone, and mutants vs WT. *p < 0.05, **p < 0.001, ***p < 0.0001. For values and number of molecules of all conditions measured, see Extended Data Table 1. Mean and s.e.m. of *k*_*cat*_ s are from two protein preparations with 5-7 replicates for each condition (see Extended Data Fig. 8 and methods).

| | $k_0$ (s<sup>-1</sup>) | $\delta$ (nm) | $k_{cat}$ (s<sup>-1</sup>) | $k_{attach}$ (s<sup>-1</sup>) |
| --- | --- | --- | --- | --- |
| **WT** | 102 ± 4 | 1.31 ± 0.03 | 4.5 ± 0.4 | 4.7 ± 0.5 |
| **+dATP** | 168 ± 7 *** | 1.49 ± 0.10 | 5.2 ± 0.4 * | 5.4 ± 0.4 |
| **+DMSO** | 87 ± 4 ** | 1.24 ± 0.04 | 3.8 ± 0.1 | 4.0 ± 0.1 |
| **+A2876 (1)** | 87 ± 2 | 1.45 ± 0.13 | 3.7 ± 0.1 | 3.8 ± 0.1 |
| **+C2981 (2)** | 23 ± 1 *** | 0.79 ± 0.05 *** | 4.6 ± 0.1 * | 5.7 ± 0.2 |
| **+D3390 (3)** | 55 ± 3 *** | 0.93 ± 0.03 *** | 5.1 ± 0.1 * | 5.6 ± 0.1 |
| **+OM** | 12 ± 1 *** | 0.26 ± 0.05 *** | 1.6 ± 0.1 * | 1.9 ± 0.1 |
| **+F3345 (4)** | 36 ± 2 *** | 1.27 ± 0.06 | 1.7 ± 0.1 * | 1.8 ± 0.1 |
| **D239N** | 140 ± 3 *** | 1.40 ± 0.03 * | 6.7 ± 0.5 * | 7.0 ± 0.5 |
| **H251N** | 152 ± 7 ** | 1.53 ± 0.08 * | 5.6 ± 0.4 * | 5.8 ± 0.4 |
| **A223T** | 104 ± 4 | 1.48 ± 0.09 | 3.2 ± 0.2 * | 3.3 ± 0.3 |
| **R237W** | 155 ± 15 * | 1.25 ± 0.06 | 2.2 ± 0.4 * | 2.2 ± 0.4 |
| **S532P** | 80 ± 8 * | 1.37 ± 0.06 | 1.4 ± 0.1 * | 1.4 ± 0.1 |

**Figure 3:**
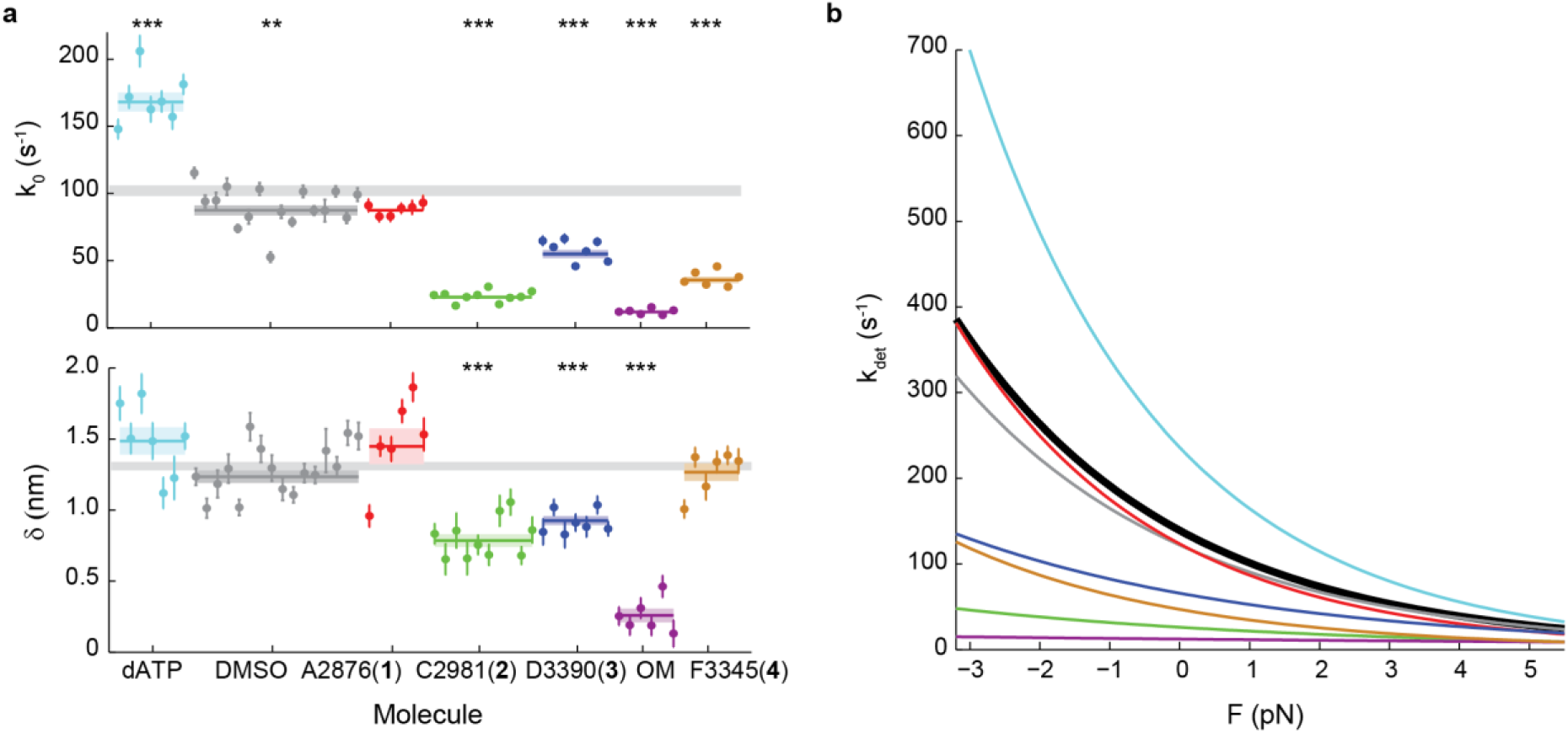
Effects of small molecule compounds on the load-dependent kinetics of single molecules of β-cardiac myosin. **a**, *k*_*0*_ (top) and *δ* (bottom) for all molecules measured. Note that some molecules have error bars smaller than the displayed data point. Weighted means ± s.e.m. are shown as horizontal lines and shaded areas, with those of untreated myosin shown as gray bars (replicated from Fig. 1) across the plots. In the case of 2-deoxy-ATP (dATP), 2 mM dATP was used in place of ATP. All other conditions used 2 mM ATP, 2% DMSO, and 25 μM compound concentration. 2-tailed unequal variances t-test was performed on dATP vs ATP, DMSO vs WT, and compounds vs DMSO. **p<0.001, ***p<0.0001. For detailed values, see Table 1 and Extended Data Table 1. **b**, The exponential dependence of detachment rate *k*_*det*_on force for every compound. Curves correspond to the weighted means of *k*_*0*_ and *δ* (see Extended Data Figs. 2 and 3 for curves of individual molecules). Myosin without treatment is shown in black. The characteristic load-dependent kinetics curve of cardiac myosin is modulated to a gradient of degrees on the single molecule level by the compounds.

### Cardiomyopathy-causing mutations alter the load-dependent kinetics of single molecules of human *β*-cardiac myosin

We next measured the effects of various cardiomyopathy-causing mutations on the load-dependent detachment rates of single molecules of human *β*-cardiac myosin using HFS (Fig. 4 and Extended Data Figs. 2 and 4, Table 1 and Extended Data Table 1). Two early-onset HCM mutations (D239N and H251N) were found previously to cause myosin to have higher actin sliding velocities in the in vitro motility assay ^32^. Consistent with their hyperactivities, these two HCM mutants exhibited faster detachment rates than the wild-type (WT) protein at the single-molecule level (D239N: *k*_*0*_ = 140 ± 3 s^−1^, p = 0.04; H251N: 152 ± 7 s^−1^, p = 0.02) and had minimal effects on force sensitivity (D239N: *δ* = 1.40 ± 0.03 nm, p = 0.04; H251N: *δ* = 1.53 ± 0.08 nm, p = 0.02). Of the three DCM mutations we studied, A223T did not change the load-dependent detachment rate compared to WT myosin (*k*_*0*_ = 104 ± 4 s^−1^, p = 0.17; *δ* = 1.48 ± 0.09 nm, p = 0.13), R237W increased the rate without changing force sensitivity (*k*_*0*_ = 155 ± 15 s^−1^, p = 0.004; *δ* = 1.25 ± 0.06 nm, p = 0.43), and S532P decreased the rate slightly without changing force sensitivity (*k*_*0*_ = 80 ± 8 s^−1^, p = 0.02; *δ* = 1.37 ± 0.06 nm, p = 0.57). Thus, point mutations that cause disease can indeed alter the load-dependent kinetics of cardiac myosin to various degrees, but whether the mutation is clinically HCM or DCM does not dictate the direction of the change (Extended Data Fig. 9). Interestingly, the DCM mutants R237W and S532P both had greater molecule-to-molecule variabilities in *k*_*0*_ compared to WT and other mutants (Fig. 4a and Extended Data Fig. 4), suggesting that these mutations may cause a disruption in protein folding that affects individual myosin molecules in a population to different extents.

**Figure 4:**
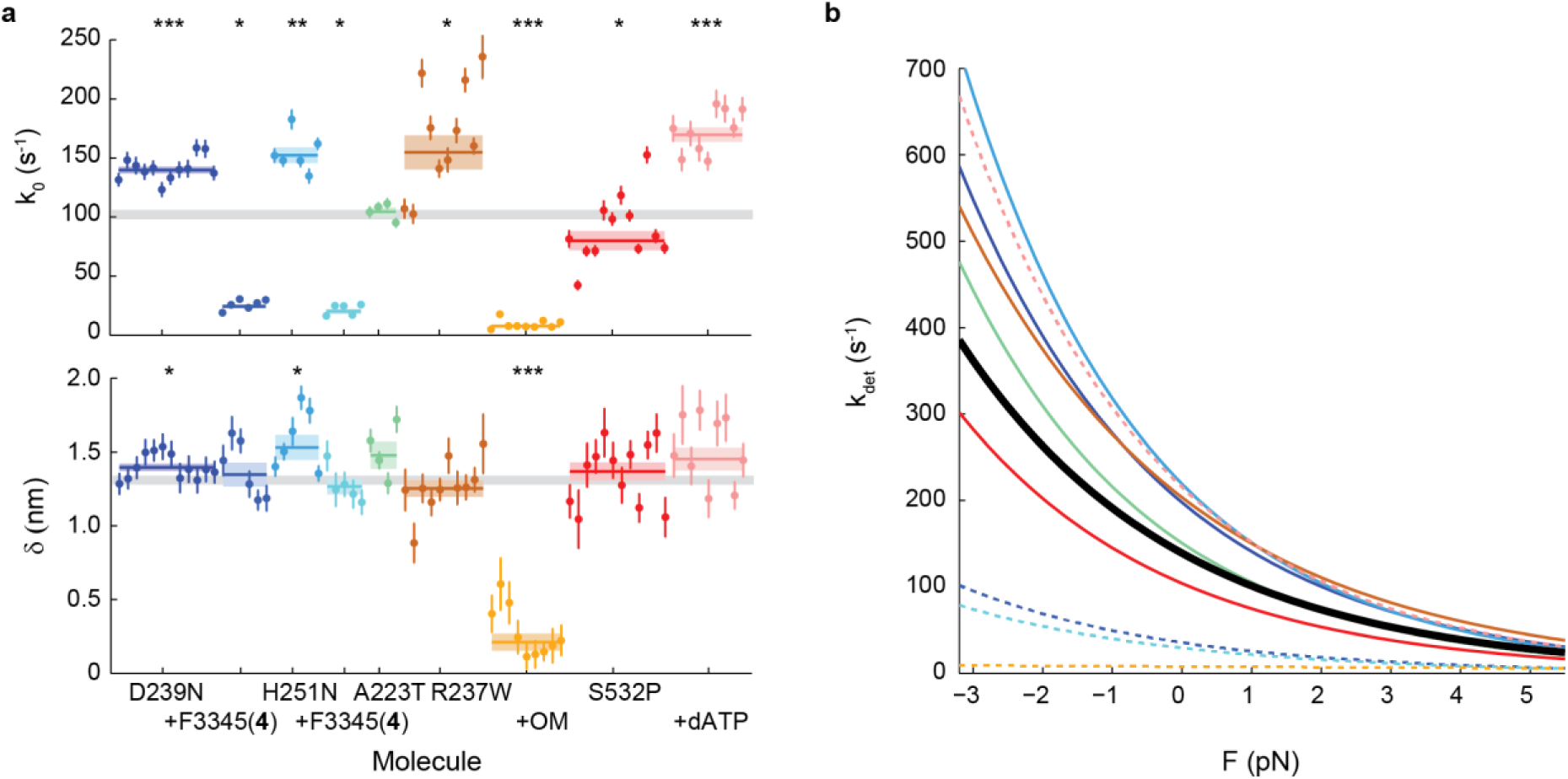
Effects of cardiomyopathy-causing mutations on the load-dependent kinetics of single molecules of β-cardiac myosin, and their reversal by small molecule compounds. **a**, *kc* (top) and *δ* (bottom) for all molecules measured. Note that some molecules have error bars smaller than the displayed data point. Weighted means ± s.e.m. are shown as horizontal lines and shaded areas, with those of untreated WT myosin shown as gray bars (replicated from Fig. 1) across the plots. Compound F3345 (4), a cardiac myosin inhibitor, was used on the HCM mutants D239N and H251. No treatment was applied to the DCM mutant A223T which showed no significant change from WT. The cardiac myosin activator OM was used on the DCM mutant R237W. Addition of allosteric compounds were always in presence of 2% DMSO. Treatment for the DCM mutant S532P was using 2 mM dATP in place of ATP. 2-tailed unequal variances t-test was performed on mutants vs WT and mutants + compounds + DMSO vs mutants + DMSO, or S532P + ATP vs S532P + dATP. *p<0.05, **p<0.001, ***p<0.0001. For detailed values, see Table 1 and Extended Data Table 1. **b**, The exponential dependence of detachment rate *k*_*det*_ on force for every mutant. Curves correspond to the weighted means of *kc* and *δ* (see Extended Data Figs. 2 and 4 for curves of individual molecules). WT is shown in black. Dashed lines are mutants with treatment. Mutations modulate the load-dependent kinetics curve of cardiac myosin on the single molecule level to varying degrees, and their effects can be reversed by treatment with compounds.

### The effects of cardiomyopathy-causing mutations can be reversed by appropriate small molecule compounds

If an effect of a disease-causing mutation is to alter the detachment rate, then we hypothesized that a small molecule compound with the opposite effect on WT myosin will reverse the alteration caused by the mutation, provided the mutation does not interfere with the action of the compound. To test our hypothesis, we first measured the effects of the inhibitor F3345 (4) on the load-dependent kinetics of each of the two HCM mutants. As expected, we observed a dramatic decrease in the detachment rates of both HCM mutants (D239N + F3345 (4): *k*_*0*_ = 24 ± 2 s^−1^, H251N + F3345 (4): 20 ± 2 s^−1^) without a significant change in force sensitivity (D239N + F3345 (4): *δ* = 1.35 ± 0.08 nm, p = 0.71; H251N + F3345 (4): 1.27 ± 0.05 nm, p = 0.17) (Fig. 4 and Extended Data Fig. 4, Extended Data Table 1). Next, since the DCM mutant R237W has a faster detachment rate than WT, we applied the activator OM and found a similarly dramatic reversal (*k*_*0*_ = 8 ± 1 s^−1^), this time with a much lower force sensitivity (*δ* = 0.21 ± 0.06 nm) as expected from the effects of OM (Fig. 4 and Extended Data Fig. 4, Extended Data Table 1). Finally, since S532P had a lower detachment rate than WT, we found that replacement of ATP with dATP reversed the mutation’s effect (*k*_*0*_ = 170 ± 6 s^−1^). Since saturating compound concentrations (25 μM allosteric effectors or 2 mM dATP) reversed the effects of mutations beyond WT levels (Fig. 4), we expect by kinetic reasoning that appropriate lower concentrations can adjust the detachment kinetics of mutants to match that of WT myosin if desired.

To this end, we then investigated the dose-dependent effects of OM on single cardiac myosin molecules’ detachment kinetics.

### Dose-dependent effects of omecamtiv mecarbil on single myosin’s detachment kinetics

Plasma concentrations of OM in patients in clinical trials are ~500 nM^27^. Therefore, we used HFS to measure the bound times of myosin incubated with OM over a range of concentrations from 30 nM to 25 μM. We found that in the presence of OM, the bound times within a force bin follow a double exponential distribution (Fig 5ab) instead of a single exponential distribution as observed under all other conditions (Fig. 1c, Extended Data Fig. 5). The presence of two populations suggests that for each event, myosin either has or does not have OM bound; the kinetics of OM binding and unbinding myosin are slow such that a single exponential distribution with an average rate is not observed^33^. We propose the following model (Fig 5d): a fraction *A* of events do not have OM bound to myosin and thus undergo detachment from actin via rate *k*_*1*_, which is presumably the same rate *k*_*det*_(*F*) measured for WT myosin in the absence of compounds. In the fraction *1-A* of events which do have OM bound, OM must first unbind myosin at rate *k*-_*OM*_ before the rest of the actin-bound state proceeds at rate *k*_*1*_. This model is consistent with the structural finding that the OM binding pocket is incompatible with the post-stroke state^34^.

**Figure 5.**
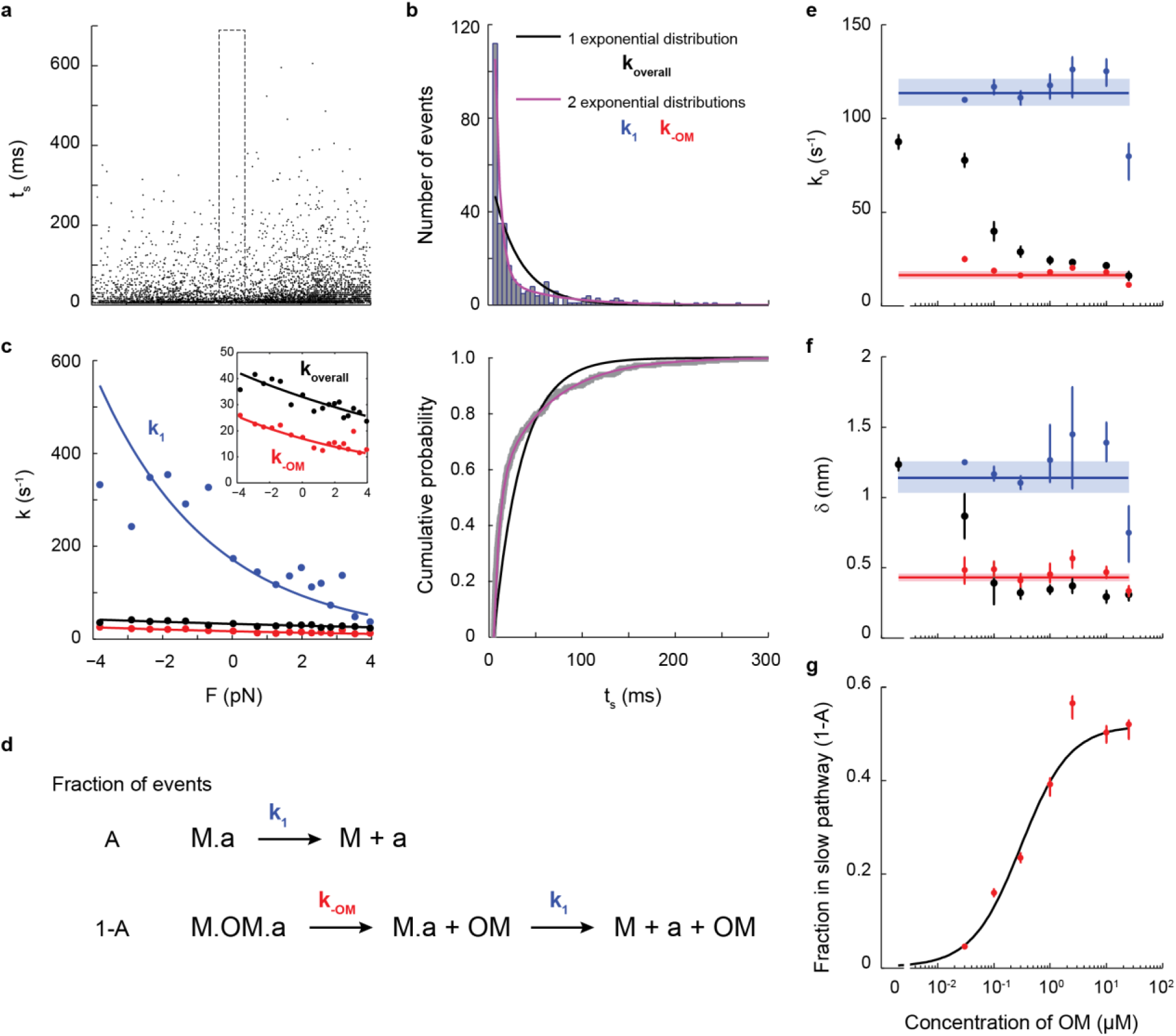
Dosage analysis of OM reveals its mechanism of action on single molecules of human β-cardiac myosin. **a**, Lifetimes of all events (N=5424) from 7 myosin molecules measured at 300 nM OM. **b**, (top) Within each force bin (the F=0 force bin is shown, as outlined in a), the binding times follow a double exponential distribution (purple) in which the fast *k*_*1*_ and slow *k*-_*OM*_rates describe the kinetics of steps illustrated in d. The rates were obtained by a MLE using MEMLET^35^. The comparison between single vs. double exponential fits is also shown in a plot of cumulative probability (bottom). **c**, Rate as a function of force (Eqn. 1) as determined from all events of the 300 nM OM dataset. The overall rate (black) was obtained by fitting a single exponential distribution. When the double exponential distribution is used, the two fitted rates *k*_*1*_ (blue) and *k*-_*OM*_ (red) each follow Eqn. 1. The curves were obtained from a MLE using MEMLET on all events without binning by force, which produced fit parameters *k*_*0*_ and *δ* in the case of the single exponential distribution, and *k*_*0&#x005F;*_, *δ*_*1*_, *k*_*0_oM*_, *δ*_*om*_, and *A* (the fraction of events in the no-OM pathway; see d) in the case of the double exponential distribution. Independent from the MLE on all events without binning by force, the rates at each force were obtained by a MLE on events binned by force (as in b) and plotted as data points. Inset shows a zoomed-in view. **d**, Simple kinetic model of OM’s action on single myosin molecules. The fraction *A* of events in which OM is not bound to myosin (“M”) undergo detachment from actin (“a”) at rate*k*_*1*_, the load-dependent detachment rate of WT myosin as measured earlier in this work (top). In the fraction *1-A* of events in which OM is bound to myosin, OM unbinds from myosin first at rate *k_-OM_,* and then myosin detaches from actin at rate *k_1_.* e-f, *k*_*0*_ and *δ* at different concentrations of OM for the overall detachment (black) and its fast (blue) and slow (red) components, obtained by MLE using MEMLET on all events from 6-9 molecules for each concentration of OM (as in c). Weighted means ± s.e.m. are shown as horizontal lines and shaded areas. Their values are: *k*_*0_1*_ = 114 ± 6 s^−1^, *δ*_*1*_ 1.14 ± 0.09 nm, *ko_OM* = 16 ± 2 s^−1^, & *δ*_*om*_ = 0.43 ± 0.03 nm. g, The fraction *1-A* of events with OM bound at different concentrations of OM. A fit to the Michaelis-Menton equation gives an apparent *K*_*d*_ = 0.31 ± 0.05 μM and *max*(*1-A*) = 0.53 ± 0.04. Error bars of the fitted parameters *k_0_1_, δ_1_, k_0_oM_,δ*_om_, and *A* in e-g are the 68% confidence interval from bootstrapping 1000 iterations using MEMLET.

Event lifetimes are MLE-fitted using MEMLET^35^ to a double exponential distribution whose two characteristic rates *k*_*1*_ and *k*-_*OM*_ follow Eqn. 1 independently (see Methods for details of the distribution used). The fitted variables are *k*_*0_1*_, *δ*_*1*_, *k*_*0_oM*_, *δ*_*om*_, and *A*. The analysis process is illustrated for the 300 nM OM case in Fig. 5a-c. As expected, whereas the overall rate *k*_*overall*_from fitting to a single exponential distribution has *k*_*0*_ and *δ* that decrease as OM concentration increases (black in Fig. 5e-f), the parameters describing each population (*k*_*0_1*_, δ_*1*_, *ko_oM*, and *Som*) are independent of OM concentration (Fig. 5e-f). The values of *k*_*0_1*_ and *δ*_*1*_(*k*_*0_1*_ = 114 ± 6 s^−1^, *δ*_*1*_= 1.14 ± 0.09 nm) are consistent with the detachment kinetics of WT myosin from actin (Fig. 1e, Table 1). The values of *ko_OM* and *Som*(*k*_*0_OM*_ = 16 ± 2 s^−1^, *δ*_*om*_ = 0.43 ± 0.03 nm) are interpreted as the detachment kinetics of OM from myosin. The non-zero positive *Som* suggests that the unbinding of OM is also slightly force-sensitive: a force in the direction of the powerstroke likely destabilizes OM’s binding pocket^34^, speeding up its release.

The fraction of OM-bound events (1*-A*) increases with OM concentration (Fig. 5g). A fit to the Michaelis-Menton equation gives an apparent *K*_*d*_ = 0.31 ± 0.05 μM, in perfect agreement with the affinity of OM to myosin’s pre-powerstroke state measured by isothermal titration calorimetry^34^. Interestingly, this fraction (*1-A*) does not reach 1 even at saturating OM concentrations; ~50% of events are able to escape the slow pathway involving OM (Fig. 5g and Extended Data Fig. 5). This result suggests that there must be a state in which myosin either does not bind the drug or binds it in a different manner such that its release is fast.

The above dosage analysis demonstrates that OM modulates the load-dependent kinetics of cardiac myosin via the same mechanism at both high and low, clinically-relevant concentrations. Taken together, we have found that both small molecule effectors and disease-causing mutations in human β-cardiac myosin can independently and in combination modulate the behavior of single myosin molecules under load. This allows one to have a fine-tuned control over the fundamental load-dependent kinetics curve that defines cardiac myosin.

### Single molecule load-dependent kinetics is the basis of the ensemble force-velocity relationship in β-cardiac myosin

In ensemble motility, the sliding velocity of an actin filament is limited by how fast the attached myosin molecule can let go of it, i.e., the detachment rate of myosin^36^. Given enough heads on the surface and a relatively short tether, as is the case in our motility assays with cardiac sS1, velocities are detachment-limited rather than attachment-limited^37^. Therefore, here, ensemble velocities measured by the in vitro motility assay should be proportional to the single molecule detachment rates measured by the optical trap. As expected, ensemble velocities were found to be linearly related to the detachment rate *k*_*0*_ with a slope of 8.9 nm and R^2^ = 0.81 (Fig. 6a). In contrast, linear regression of velocity vs. attachment rate *k*_*attach*_ (see Discussion for calculation of *k*_*attach*_ by Eqn. 3) gave R^2^ = 0.35 (Extended Data Fig. 6). In simplest terms, since unloaded velocity *v = d/t_s_ = d*k_0_,* where *d* = step size and *t*_*s*_ = strongly bound state time, the slope obtained from the linear fit between velocity and *k*_*0*_ can be interpreted as the effective step size of myosin in the ensemble. This effective step size (8.9 nm) is greater than the value from direct step size measurements on single cardiac myosin molecules (~7 nm) ^10,38,39^ because the non-zero load-dependence *(δ)* of cardiac myosin results in faster detachment rates in the ensemble ^40^. Interestingly, the activators and HCM mutants lie on or above the fitted line while the inhibitor F3345 (4) and DCM mutants lie below it (Fig. 6a). This presents a possible distinction between the two groups in which activating perturbations produce greater coordinated movement in the ensemble than predicted by single molecule detachment rates while inhibitory perturbations result in less enhancement.

**Figure 6:**
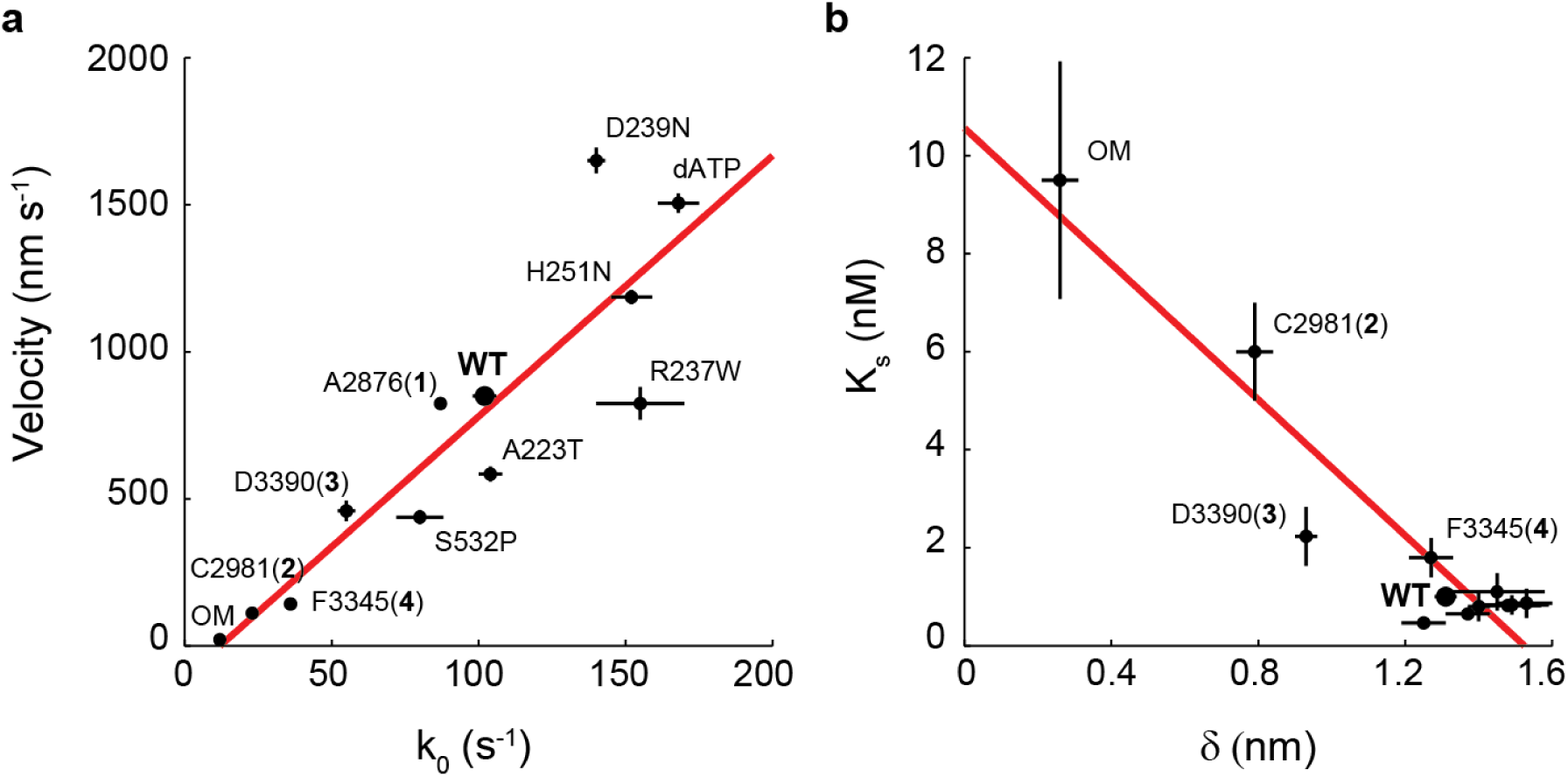
Single molecule load-dependent kinetics as the basis of the ensemble force-velocity relationship in β-cardiac myosin. **a**, Unloaded actin sliding velocity measured by in vitro motility assay vs. *k*_*0*_, the detachment rate at zero load for a single molecule of myosin, for the compounds and mutations studied. The rate of detachment limits ensemble velocity. Linear regression gives a slope of 8.9 nm and R^2^ = 0.81. Note that some points have error bars smaller than the dot. **b**, The parameter *K*_*s*_ from the loaded motility assay is a measure of load-bearing ability at the ensemble level, while *δ* is the load-sensitivity parameter determined at the single molecule level. Pearson correlation −0.94. For clarity, the cluster of data points at the lower right corner are not individually labeled. Motility velocity and *K*_*s*_ values of HCM, DCM, and dATP are obtained from other studies ^20,25,32^ (Ujfalusi Z., Vera C., Mijailovich S., Svicevic M., Choe Yu E., Kawana M., Ruppel K., Spudich J., Geeves M., and Leinwand L., *JBC* manuscript under review; Tomasic I., Liu C., Rodriguez H., Spudich J.A., Bartholomew Ingle S.R., manuscript in preparation).

In addition to *k*_*0*_, our data revealed differences in force sensitivity *(δ)* under different perturbations to cardiac myosin at the single molecule level (Figs. 3a, 4a). Therefore, we asked whether changes to force sensitivity of single molecules result in similar findings on the ensemble level. We measured ensemble force sensitivity by a loaded in vitro motility assay ^20^. In this assay, utrophin on the surface binds to actin to provide the load against which myosin must overcome in moving actin filaments. Increasing concentrations of utrophin increase load and thus decreases actin sliding velocity. This gives an ensemble force-velocity curve (Extended Data Fig. 7). The concentration of utrophin *K*_*s*_ required to reduce velocity by half is therefore a measure of force sensitivity. As expected, we found that *K*_*s*_ is inversely proportional to *δ*(Pearson correlation coefficient = −0.94) (Fig. 6b and Extended Data Fig. 7). As single myosin molecules become insensitive to load (smaller *δ*), such as the cases of the activators OM, C2981 (2), and D3390 (3), higher concentrations of utrophin are required to reduce the ensemble velocities by a half.

**Figure 7:**
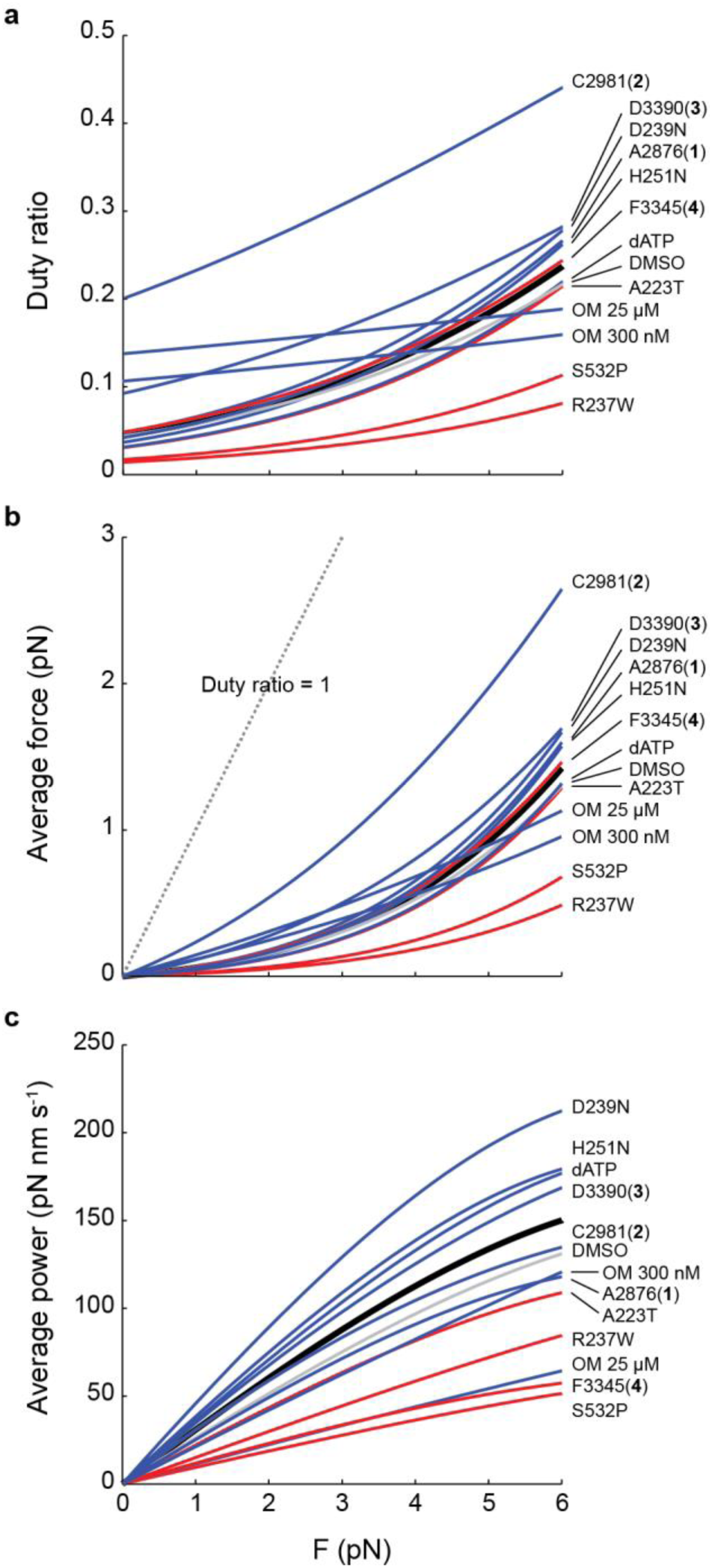
The effects of small molecule compounds and cardiomyopathy-causing mutations on the load-dependent power production of single molecules of β-cardiac myosin. **a**, Duty ratio, b, average force, and c, average power as a function of the load force *F,* calculated by Eqns. 2-5. All curves are calculated using values of detachment rate kinetics (*k*_*0*_,*δ*), *k*_*cat*_, and *k*_*attach*_ given in Table 1. Black: WT. Gray: DMSO. Blue: HCM and activators. Red: DCM and inhibitor.

In conclusion, we have found that the load-dependent kinetics inherent to each single molecule of cardiac myosin can be modulated by various small molecule compounds and mutations. This single molecule characteristic underlies the force-velocity relationship exhibited in ensemble.

## Discussion

Large sequence differences between various types of myosin lead to dramatically different scales of the load-dependent detachment rate. Myosin I ^15,16^, V ^14^, smooth ^13^, and β-cardiac ^10,11^ isoforms have distinct rates and force sensitivities ranging from *k*_*0*_ ~ 1 to 100 s^−1^, and *δ* ~ 1 to 12 nm. Here, armed with the efficient HFS method, we explored whether changes to the load-dependent rate can be made via small perturbations in the form of small molecule compounds and cardiomyopathy-causing single missense mutations in human β-cardiac myosin. The results lie across the spectrum: not all perturbations changed the single molecule load-dependent detachment rates (activator A2876 (1), DCM mutant A223T), while the rest modulated the curve to various extents (Figs. 3, 4, and Extended Data Fig. 9). Modulation was seen in the rate at zero load *k*_*0*_ (~ 10 to 170 s^−1^) and/or the force sensitivity *δ* (~ 0.2 to 1.5 nm). Furthermore, the effects of mutations could be reversed by an appropriate compound (Fig. 4). Thus we have observed fine tuning of the load-dependent kinetics on single molecules of cardiac myosin. In oversimplified terms, we can control cardiac contractility at the single molecule level.

### Perturbations to myosin change its load-dependent detachment rates through different steps in the ATPase cycle

Any step in the strongly-bound state may be affected by the compounds and mutations. ADP release is the rate-limiting step of the strongly-bound state under unperturbed circumstances for WT cardiac myosin. This is likely the acted-upon step for dATP, inhibitor F3345 (4), and mutations since the transition state distance *(δ)* for those conditions remained close to that of WT myosin without treatment (Figs. 3a, 4a, and Table 1). Furthermore, it has been measured by stopped flow experiments of bovine cardiac HMM that the faster nucleotide release rate when dATP is used in place of ATP matches the detachment rate measured on single molecules (Tomasic I., Liu C., Rodriguez H., Spudich J.A., Bartholomew Ingle S.R., manuscript in preparation). For the activators C2981 (2), D3390 (3), and OM, the significantly smaller *δ* (Fig. 3a and Table 1) may suggest a new rate limiting step with a different transition state. Alternatively, the compounds may have simply changed the transition state of ADP release to yield smaller distance. Future kinetic studies are needed to distinguish between these models and illuminate the precise steps at which the compounds act.

For OM, it has been shown that the drug stabilizes the pre-powerstroke state in which myosin is ready to bind to actin ^34^, accelerates phosphate (Pi) release ^23,28,30^, then delays the power stroke while myosin is in the actin-bound state ^30^. Structural data suggests OM unbinds before the stroke ^34^, thus leaving the ADP release rate unaffected ^23,28-30^. In our study, we measured an overall large increase in the lifetime (1*/k_det_*) of the bound state, consistent with the previously observed delay in powerstroke. Through dosage analysis, we determined that this delay can be attributed to the off rate of OM for events in which myosin has the drug bound at the start.

Unlike solution measurements, measurement of event lifetimes on single myosin molecules provides direct determination of the detachment rate. But at the same time, a limitation of HFS so far is an inability to measure the powerstroke. Thus, we cannot observe the exact proposed sequence of events from myosin binding to actin, delay, stroke (presumably after OM unbinds), post-stroke (as ADP remains bound), and finally unbinding from actin. It has been suggested that OM drastically reduces the stroke size of myosin (Woody, M.S. et al. Biophysical Journal, Volume 114, Issue 3, 317a - 318 abstract). The outstanding question of whether or not myosin strokes after OM unbinds can be answered using an ultrafast feedback trap that directly measures the sequence of events.

Despite the limitations of HFS in distinguishing different phases of the bound state, it can be argued that wherever in this state an effector acts, the same result is achieved: the overall bound time is changed, and it is this overall bound time that limits ensemble velocity and determines force production.

### Single molecule vs. ensemble observations

Fitting with the above statement that the overall bound time determines ensemble velocity, we found a strong linear relationship between single molecule detachment rate and ensemble velocity (Fig. 6a). A linear relationship had also been observed between the ADP release rate, which limits detachment, and the unloaded shortening velocity of skeletal muscle fibers^17,18^. Activators C2981 (2) and D3390 (3) and inhibitor F3345 (4) may act in steps of the ATPase cycle different from those of OM, but the result is the same - a prolonged strongly bound state time. This manifests itself similarly: a slowing of ensemble velocity, despite differences in the details of the mechanism. In this way, the single molecule detachment rate *k*_*0*_ does truly dictate to a large extent motility velocity for cardiac myosin.

Since small molecule effectors may prolong the strongly bound state by slowing the kinetics of different steps, it is not surprising that the distances to the transition state are also altered. While the mechanistic details may vary with different compounds or mutations, the net result is a change in *δ,* the parameter that encapsulates force sensitivity. When the force sensitivity inherent to a single molecule is changed, the result is reflected at the ensemble level (Fig. 6b). Thus we find that different mechanisms of changing *k*_*0*_ and *δ* can lead to the same ensemble force-velocity behavior (Fig. 6).

While the ensemble observation correlates well with predictions from single molecule data, not all the variation observed in the ensemble case can be explained by the current single molecule model (Fig. 6). For example, why do HCM mutations and activators move faster in motility than predicted by detachment rates alone, while DCM mutations and the inhibitor F3345 (4) have the opposite effect? Building upon the measured single molecule parameters, a more sophisticated model of ensemble motility may incorporate effects of attachment kinetics^37^ and can be explored in future work.

One parameter for which we have not yet measured effects of perturbations is myosin’s step size *d.* Previous studies show little or no significant change to *d* by HCM or DCM mutations ^38,39,41^, but that may not be the case for all perturbations studied in this paper (Woody, M.S. et al. Biophysical Journal, Volume 114, Issue 3, 317a - 318 abstract). Future studies will answer this question. In addition, one may ask whether myosin is truly making a stroke under an applied load. Although we cannot observe the working stroke using HFS, we do believe that myosin is able to stroke because we apply average forces below ~5-6 pN (Fig. 1c). It has been shown for skeletal muscle myosin that the stroke size remains the same at ~5 nm up to the isometric force of 5.7 pN ^12^. This was found to be true when very short events (<1 ms) corresponding to short, unproductive binding were excluded from the analysis. In HFS, the stage oscillation of 200 Hz set our lower detection limit at 5 ms (one period), so all events included in our analysis have lifetimes greater or equal to 5 ms and therefore are expected to have productive working strokes. However, direct observation of a stroke under load would require an ultrafast feedback optical trap system ^12^.

We have found that small perturbations, activating or inhibitory, can increase, decrease, or not affect the detachment rates (Figs. 3, 4, and Extended Data Fig. 9). While this may be expected given that compounds and mutations can act via different mechanisms of action, the question remains: what distinguishes one group from the other? A heart containing an HCM mutation (eg. H251N) undoubtedly is hyper-contractile ^42^ compared to one bearing a DCM mutation (eg. R237W) despite both having faster detachment rates. Armed with knowledge of their load-dependent kinetics, we believe that the answer lies in an assessment of the resulting duty ratio, average force, and power output, as explained in the following section.

### Implications of single-molecule load-dependent kinetics on power production by β-cardiac myosin

The duty ratio is the time myosin spends in the strongly bound state *t*_*s*_ divided by its entire cycle *t*_*c*_ and is an important parameter because it represents the fraction of time myosin is in a force producing state. The entire cycle is made up of time in the strong and weak states, *t*_*s*_ and *t_w_,*respectively. The duty ratio as a function of load force *F* can be expressed as

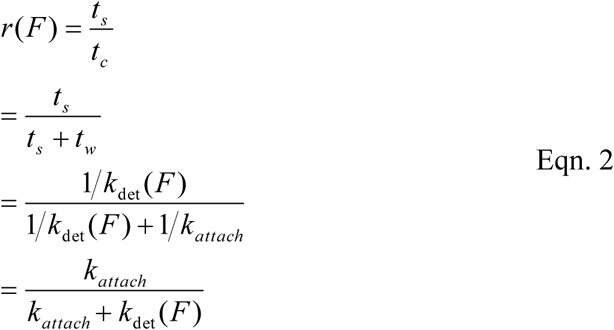

where *k*_*attach*_ describes the lifetime of the weak binding state and is assumed to be independent of force since myosin cannot yet sense any load on actin when unbound. *k*_*attach*_ cannot be determined by directly measuring the lifetimes of the unbound state in the optical trap because there, the rate of attachment is diffusion-limited via the positions of the actin dumbbell and myosin. However, *k*_*attach*_ can be calculated by

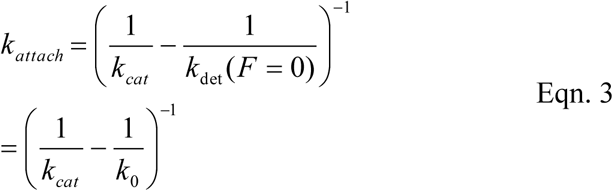

where *k*_*cat*_ is the overall cycle rate measured by steady state ATPase. We know the values of all variables in these equations: *k*_*det*_(*F*) given by Eqn. 1 is measured by HFS in this study, *k*_*cat*_ is measured in an actin activated ATPase assay in this study (Table 1 and Extended Data Fig. 8) and others ^20,32^ (Ujfalusi Z., Vera C., Mijailovich S., Svicevic M., Choe Yu E., Kawana M., Ruppel K., Spudich J., Geeves M., and Leinwand L., *JBC* manuscript under review). Therefore we can calculate the duty ratio as a function of load force (Fig. 7a). Values of *k*_*0*_, *δ, k*_*cat*_, and *k*_*attach*_ used are given in Table 1. Interestingly, we find that the duty ratios of the activators and HCM mutations in general lie above those of the inhibitor and DCM mutations. This result is consistent with physiological expectations that HCM mutations and activators cause myosin to have a higher duty ratio, while DCM mutations and inhibitors result in the opposite^28-30,34,43^.

**Figure 8:**
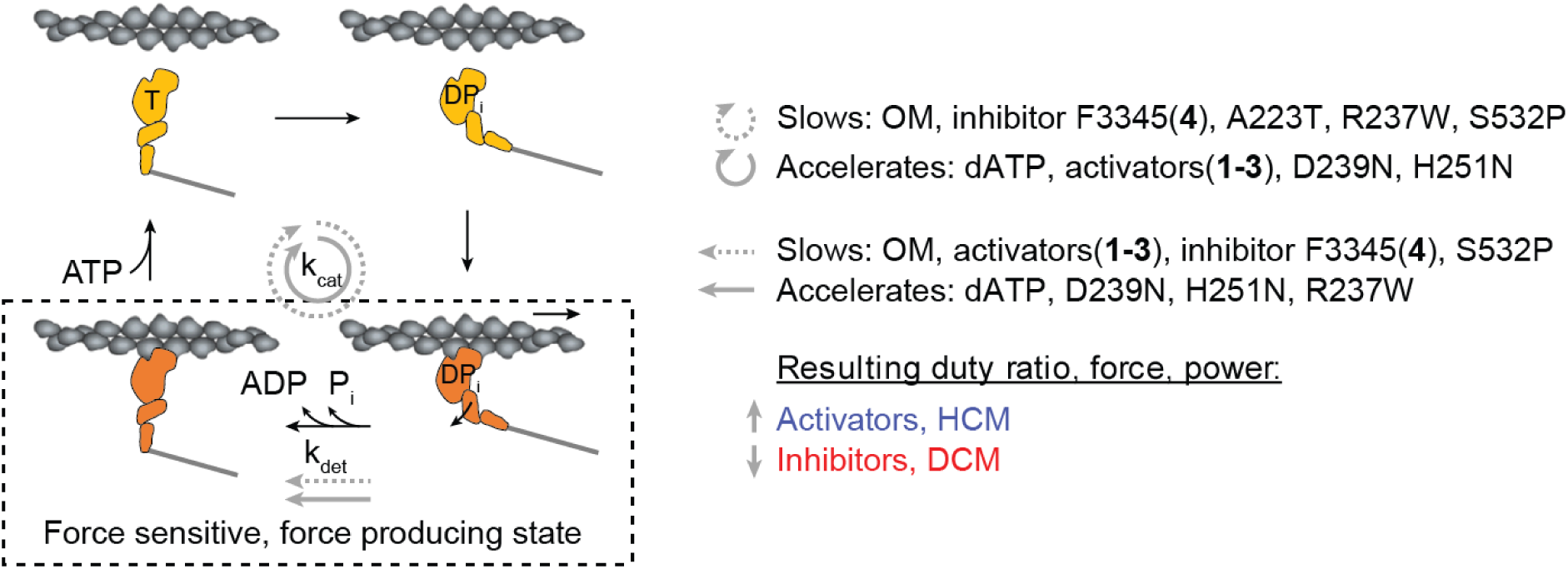
Model of the effects of small molecule compounds and cardiomyopathy-causing mutations on the contractility of β-cardiac myosin at the single molecule level. The overall cycle, described by *k*_*cat*_, can be slowed or accelerated by activators, inhibitors, HCM, and DCM mutations. Likewise, time spent in the strongly bound, force sensitive, force producing state can also be prolonged or shortened by these perturbations. It is the aggregate effects on the parameters of power production which discriminates activating (small molecule activators, HCM mutations) from inhibitory perturbations (small molecule inhibitors, DCM mutations).

The inhibitor F3345 (4) is an outlier whose curve lies among those of the activators/HCM’s. Compared to the absence of small molecule compounds, OM dramatically increases the duty ratio of cardiac myosin even at low forces, consistent with previous studies which proposed that it causes myosin to spend a much greater fraction of time in the strongly bound state ^28-30,34^ At higher forces, its duty ratio does not increase as much as that of others due to its low force sensitivity (*δ*) (Fig. 3). Surprisingly, two of the activators (C2981 (2) and D3390 (3)) increased the duty ratio even more dramatically than OM at both high and low forces. Furthermore, because their force sensitivities were not as low as that of OM (Fig. 3), their duty ratios increased more steeply with increasing force (Fig. 7a).

Average force is the load force *F* against which a single molecule was able to stroke multiplied by the duty ratio:

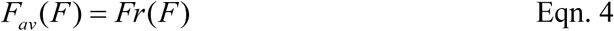

The average force myosin produces increases with load force until the isometric force, taken to be 6 pN here ^12^ (Fig. 7b). As expected, the activators C2981 (2), D3390 (3) and OM increased the average force the most because they had the largest increase in duty ratio. Similar to duty ratio, calculation of the average force broadly discriminates the activators and HCM mutations from the inhibitor and DCM mutations.

Finally we calculate the average power produced by a single myosin:

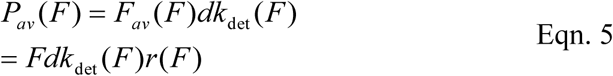

where step size *d* is taken to be 7 nm. A constant step size *d* may be a simplification as some perturbations may alter it (see previous discussion). Nevertheless, it is illustrative to calculate the power produced by single myosins to understand the effects of changes to the detachment rate. Here again, HCM and activators overall segregate from DCM mutations and inhibitors (Fig. 7c). The activator OM is the notable exception: at 25 μM, it causes decreased power despite a higher average force because of its much slower kinetics; at 300 nM, while the dramatic power reduction is alleviated, its power is still on the lower end of the group of activators. This can help explain the effects of OM observed at the organ level: the predicted higher average force is consistent with an increase in peak time-dependent elastance of the left ventricle (a measure of LV contractility) and stroke volume, while the slower kinetics is consistent with a longer time to maximum pressure and a longer time in systole ^23,27^.

Our dosage analysis of OM revealed that a common molecular mechanism underlies the drug’s effect across a large range of concentrations, including lower, clinically relevant doses (~500 nM)^27^ (Fig. 5). At the same time, the sensitivity of myosin’s power production to the concentration of OM, as discussed above (Fig. 7c), illustrates the importance of the therapeutic window: a careful balance of OM’s slow kinetics with its activating function to increase the duty ratio must be and is considered in clinical trials. Finding the optimal degree of alteration in myosin function is critical in modulating ventricular function as a whole.

The *k*_*cat*_term in the above equations should be measured in future studies on heavy meromyosin (HMM) rather than sS1 in order to encapsulate the term *N_a_,* the number of available heads. The myosin “mesa” is a relatively flat surface on the motor domain that is involved in the interaction of folded back heads with the proximal tail, forming the so-called interacting heads motif, and which harbors many HCM mutations.^44^ Mutations on the mesa and compounds which regulate the interacting heads motif may change *N*_*a*_ via the proportions of the folded back state, ^45-49^ thus changing *k*_*cat*_ measured for HMM while showing no change for sS1. In addition, other factors that build a more complex, comprehensive system (eg. regulated thin filament, myosin binding protein C) can also modulate the folded back state and thus change *k*_*cat*_. These unaccounted-for complexities may explain why the compound A2876 (1) showed no significant change from WT/DMSO in all experiments and the power calculation performed in this study using the simple purified actin-activated myosin system (Figs. 2, 3, 7, and Extended Data Fig. 8), yet was discovered as an activator from an ATPase screen using regulated thin filaments.

Given the above considerations, we propose a model in which activating vs inhibitory perturbations of cardiac myosin are discriminated by the aggregate result on duty ratio, average force, and ultimately average power (Fig. 8). While the load-dependent detachment rate is an essential parameter since it determines the time myosin spends in a strongly bound, force producing state, it alone cannot distinguish the two types of perturbations. The steady state *k*_*cat*_ cannot, by itself, distinguish them either. When both are taken into account by a calculation of the duty ratio, average force, or average power, then the distinction is made. To build a comprehensive understanding of the molecular mechanism underlying muscle contraction, these calculations takes into consideration all the parameters of force production: load force *F*, the load-dependent detachment rate *k*_*det*_(F), the overall cycle rate *k*_*cat*_, the number of functionally available heads *N*_*a*_ (included in *hat),* and the step size *d.*

Recent studies on the HCM mutations R453C ^38^, R403Q ^39^, and three in the converter domain ^50^ found variable changes to parameters of force production (without *k*_*det*_(F) measurements) which do not clearly point to a hyper-contractile phenotype. Armed with the efficiency of the HFS method, future studies measuring *k*_*det*_*(F)* for these mutants and others will illuminate whether molecular hyper-contractility and hypo-contractility are indeed the respective underlying effects of HCM and DCM.

## Methods

**Small molecule compounds** were a generous gift from MyoKardia Inc. (South San Francisco). They are stored at 10 mM in DMSO at 4 °C.

### Protein preparation

Construction, expression and purification of the WT and mutated recombinant human β-cardiac myosin sS1s are described in detail elsewhere ^32,38^. Briefly, a truncated version of *MYH7* (residues 1 - 808), corresponding to sS1, with either a C-terminal enhanced green fluorescent protein (eGFP) or a C-terminal eight-residue (RGSIDTWV) PDZ-binding peptide was co-expressed with myosin light chain 3 *(MYL3)* encoding human ventricular essential light chain (ELC) and containing an N-terminal FLAG tag (DYKDDDDK) and tobacco etch virus (TEV) protease site in mouse myoblast C2C12 cells using the AdEasy Vector System (Obiogene Inc.). The myosin heavy chain with its associated FLAG-tagged ELC was first purified from clarified lysate with anti-FLAG resin (Sigma). After cleaving off the FLAG tag with TEV protease, the human β-cardiac sS1 was further purified using anion exchange chromatography on a 1 mL HiTrap Q HP column (GE Healthcare). Peak fractions were eluted with column buffer (10 mM Imidazole, pH 7.5, ~200 mM NaCl, 4 mM MgCh, 1 mM DTT, 2 mM ATP and 10% sucrose) and concentrated by centrifugation in Amicon Ultra-0.5 10 kDa cutoff spin filters (EMD Millipore) before being stored at −80°C. The purity of the protein was confirmed using SDS-PAGE (Extended Data Fig. 1). Frozen proteins exhibited similar activities in the motility, single-molecule, and ATPase assays compared to their fresh counterparts, as seen in previous work ^50^.

### Deadheading myosin

Myosin protein used for motility and trap experiments was further subjected to a “deadheading” procedure to remove inactive heads. Myosin was mixed with 10x excess of unlabeled F-actin on ice for 5 min, followed by addition of 2 mM ATP for 3 min, then centrifuged at 95K rpm in a TLA-100 rotor (Beckman Coulter) at 4 °C for 20 min. The supernatant was collected to be used in the motility and trap experiments. Myosin used in the ATPase assay was not subjected to deadheading.

### Actin-activated ATPase

Purified bovine cardiac G-actin provided by MyoKardia Inc. was freshly cycled to F-actin by extensive (4 times over 4 days) dialysis in ATPase buffer (10 mM Imidazole, pH 7.5, 5 mM KCl, 3 mM MgCl2 and 1 mM DTT) to remove any residual ATP up to a week before each experiment. The monomeric concentration of F-actin was determined by measuring the absorbance of a serial dilution of the actin in 6 M guanidine hydrochloride both at 290 nm with an extinction coefficient of 26,600 M^−1^ cm^−1^ and at 280 nm with an extinction coefficient of 45,840 M^−1^ cm^−1^ in a spectrophotometer (NanoDrop). Full-length human gelsolin was added to actin at a ratio of 1:1000 to reduce the viscosity of the actin and thereby decrease pipetting error at higher actin concentrations without affecting the ATPase activity ^38^. The steady-state actin-activated ATPase activities of freshly prepared human β-cardiac sS1-eGFP were determined using a colorimetric readout of phosphate production ^51^. In this assay, reactions containing sS1 at a final concentration of 0.01 mg mL^−1^, 2 mM ATP or dATP, and actin at concentrations ranging from 2 to 100 μM were performed at 23°C with plate shaking using a microplate spectrophotometer (Thermo Scientific Multiskan GO). Prior to the start of each reaction, sS1s were incubated on ice with 25 μM of the indicated type of small molecule compound in 2% DMSO for 5 min. ATP or dATP was added at t = 0, and four additional time points up to 30 min were measured subsequently for each actin concentration. The rate of sS1 activity was obtained by linear fitting the phosphate signal as a function of time and converted to activity units using a phosphate standard.

A set of experiments measuring all conditions (dATP and compounds) on WT sS1-eGFP were performed on the same day that myosin was purified to ensure that conditions can be directly compared to each other without variation due to different protein preparations. Each experiment of each condition was performed in duplicates or triplicates. The error on each data point (ATPase activity at a certain actin concentration) represents the s.e.m. of the replicates. The Michaelis-Menten equation was fitted to determine the maximal activity (*k*_*cat*_) and the actin concentration at half-maximum (apparent *K*_*m*_for actin). Data from one day is shown in Extended Data Fig. 8a. The *k*_*cat*_ values from each day’s experiment are averaged and presented in Table 1 along with their s.e.m.’s. Here we performed two sets of experiments using two protein preparations and a total of 5-7 replicates. Since each set of experiments measured all conditions on the same day using the same protein prep, we used the paired t-test to test for significance (Table 1). We also performed a Michaelis-Menten fit to the aggregated data from both days (Extended Data Fig. 8b).

### In vitro motility

The basic method followed our previously described motility assay with some modifications ^31,39,50^. Dose-dependent change in unloaded velocity was observed by measuring actin velocity at 8-12 different concentration of each compound. Four different concentrations of a compound were tested on each motility slide, and 2-3 slides were simultaneously analyzed in each set of experiment to obtain full dose-titration curve. Normalization of velocity was done using the velocity of the WT control from the same set of experiments. For loaded in vitro motility assays, 2-4 different concentrations for each compound (including baseline control) were chosen according to the dose-titration curve of unloaded velocity. For each compound concentration, the force-velocity curve was obtained by measuring actin velocity at 8-12 different concentrations of utrophin. Four different concentrations of utrophin were tested on each motility slide, and 2-3 slides were simultaneously analyzed in each set of experiments to obtain full force-velocity curves. Movies were analyzed using FAST (Fast Automated Spud Trekker), software that automates tracking of actin filaments for in vitro motility assays and analyzes the velocity of each filament in high throughput^20^. Velocities reported are mean velocity (MVEL) with 20% tolerance, as described previously ^20^. Each experiment tracked ~400 to ~1900 actin filament movements, and the same condition was repeated. For unloaded velocity, mean +/- standard error of mean from the two experiments are plotted against compound concentration (Fig. 2). After myosin was attached to the motility surface and washed with BSA-containing assay buffer, a final solution that contained fluorescently-labeled bovine actin, 2 mM ATP, an oxygen-scavenging system (0.4% glucose, 0.11 mg ml^−1^ glucose oxidase, and 0.018 mg ml^−1^ catalase), an ATP regeneration system [1 mM phosphocreatine (PCR), 0.1 mg/mL creatine phosphokinase (CPK)], and a designated concentration of compound in 0.5% DMSO was added. The myosin was incubated in the final solution containing compound for at least 5 minutes before movies were taken. The amount of time the final solution was incubated in the chamber before movies were taken did not affect the velocity (data not shown). DMSO had minimal effect on velocity (2% DMSO had a 10% reduction in velocity). All experiments were performed at 23 °C.

### Single molecule measurements of load-dependent detachment rates

Measurements were performed using the Harmonic force spectroscopy (HFS) method in a dual beam optical trap ^11^.

#### Sample chamber preparation

The sample chamber was built on a glass coverslip spin-coated first with 1.6 μm-diameter silica beads (Bang Laboratories) as platforms and then with a solution of 0.1% nitrocellulose 0.1% collodion in amyl acetate. The flow channel was constructed by two double-sided tapes between a glass slide and the coverslip. Experiments were performed at 23 °C. GFP antibody (Abcam ab1218) at a final concentration of ~1 nM diluted in assay buffer (AB) (25 mM imidazole pH 7.5, 25 mM KCl, 4 mM MgCl_2_, 1 mM EGTA, and 10 mM DTT) was flowed into the channel to bind the surface for 5 min, followed by washing and blocking with 1 mg ml^−1^ BSA in AB (ABBSA) for 5 min. Dead-headed human β-cardiac myosin sS1-eGFP diluted to a final concentration of ~10-50 nM in ABBSA was flowed in next and allowed to saturate GFP antibody binding sites for 5 min. Then unbound myosin was washed out with ABBSA. Lastly, the final solution was flowed in consisting of 2 mM ATP, 1 nM TMR-phalloidin-labelled biotinylated actin (Cytoskeleton, ~1 biotin per monomer) filaments, an oxygen-scavenging system (0.4% glucose, 0.11 mg ml^−1^ glucose oxidase, and 0.018 mg ml^−1^ catalase), and 1 μm-diameter neutravidin-coated polystyrene beads (ThermoFisher) diluted in ABBSA. The chamber was sealed with vacuum grease. For experiments with 2-deoxy-ATP, 2 mM 2-deoxy-ATP was used in place of ATP. For experiments with other small molecule compounds, ABBSA contained 2% DMSO and specified concentration of compound.

#### HFS experiment

Details of the technique have been presented elsewhere ^11^. Two neutravidin beads were trapped in two trap beams. The beads were moved in ~57 nm steps in a 9×9 raster scan using acousto-optic deflectors while their displacements were recorded both in brightfield (in nm) and with quadrant photodiode (QPD) detectors (in voltage) to produce a voltage-to-nm calibration. The stiffness of each trap beam was calculated by the equipartition method and was typically ~0.07 pN nm^−1^ (right) and 0.09 pN nm^−1^ (left) in these experiments. After this calibration, an actin filament was stretched to ~5 pN pre-tension between the two trapped beads to form a “dumbbell.” We sinusoidally oscillated the piezoelectric stage at 200 Hz with amplitude set at 50 nm. While the amplitude was set to 50 nm, the stage actually oscillated at ~30 nm due to the response of the stage at 200 Hz. The dumbbell was lowered near a platform bead to check for binding to a potential myosin on the platform, as indicated by large, brief increases in the position amplitudes of the trapped beads (“robust interactions”) due to stronger association with the oscillating stage. Due to compliance, the dumbbell oscillates with amplitude ~15-25 nm during a binding event, which corresponds to a total force dF ~3-4 pN from the two traps. We typically explored ~5-10 platform beads before robust interactions were observed, suggesting that the GFP antibody and myosin concentrations used in this study resulted in sufficiently low number of properly-oriented molecules on the surface such that events observed were likely due to a single myosin. Control experiments with GFP antibody only (without myosin) revealed that any interactions between the antibody and the dumbbell are not “robust” as defined above; their amplitudes are small, and the durations are very short such that they would not pass the analysis’s detection criteria as detailed below. The positions of the trapped beads and the piezoelectric stage were recorded at 50 kHz sampling frequency. We recorded data on each molecule for ~10 min and used each slide for 1.5-2 hr. We did not observe slowing down of motor during the 10-min recording for one molecule nor for multiple molecules on one slide during a 2 hr experiment, suggesting that the 2 mM ATP was not significantly depleted despite our foregoing an ATP regeneration system.

#### HFS data analysis

The theory of the HFS method and the data processing details have been presented elsewhere ^11^. In summary, in the unattached state, the dumbbell oscillates π/2 ahead of the stage due to the fluid motion while in the attached state the dumbbell oscillates in phase and with a larger amplitude due to stronger association with the stage. Therefore, events are detected based on a simultaneous increase in oscillation amplitude above a threshold (~15 nm) and a decrease in phase below a threshold (~0.5 rad) for at least one full stage oscillation period. Several additional filtering of events are applied to minimize false positives. The final number of events for each molecule ranged from ~300-1000. For each event, the mean force due to each trap is calculated as the bead position averaged over the duration of binding multiplied by the trap stiffness. The total mean force on myosin is the sum of the two mean forces. A range of forces, both positive and negative, is automatically applied to myosin because binding can occur anywhere in the cycle of oscillation. Through this event detection algorithm, the mean force, amplitude of oscillation, and duration (*F*, *ΔF, t_s_*) of each event are obtained.

For each molecule, events are binned by force in ~1 pN increments, and a maximum likelihood estimation (MLE) of the geometric distribution (the discrete version of the exponential distribution) of bound times, with dead-time correction (5 ms), is performed for each bin to obtain the detachment rate for the mean force of the bin, *k_det_(F).* The error on *k_det_(F)* is calculated from the variance of the MLE as the inverse Fisher information. This results in a plot of *k_det_(F)* vs. *F* to which we perform a weighted least-squares fit of Eqn. 1 using *k*_*0*_ and *δ* as fitting parameters. Theoretical errors on the estimates for *k*_*0*_ and *δ* are calculated from the inverse Fisher information matrix. Thus we have *k*_*0*_ and *δ* and their errors for each molecule. Then, values of *k*_*0*_and *δ* from each molecule are averaged weighted by their errors (variance) to obtain mean values for each condition. The errors on these mean values are calculated as the s.e.m.

We have also used MEMLET^35^ to analyze a few molecules to compare the two analysis methods. The input data to MEMLET is the same (*F,ΔF*, *t*_*s*_) produced by the HFS analysis program. MEMLET does not require binning events in force. Instead, it produces the fitted variables *k*_*0*_and *δ* from a MLE fit of all events of one molecule to the geometric distribution in which the rate parameter of the distribution is given by Eqn. 1. In the molecules we tested, MEMLET gave nearly identical *k*_*0*_ and *δ* values. MEMLET uses bootstrapping to obtain the uncertainties of the fitted variables. The 68% confidence intervals obtained by MEMLET bootstrapping were very similar to the errors calculated by the inverse Fisher information matrix. The excellent agreement between these two analysis methods, one which bins in force and the other which does not, is expected because the large number of events available through the HFS method minimizes the negative effects of binning^35^.

#### Analysis of single molecule OM dosage data

According to the model in Fig. 5d, the distribution of bound times is given by

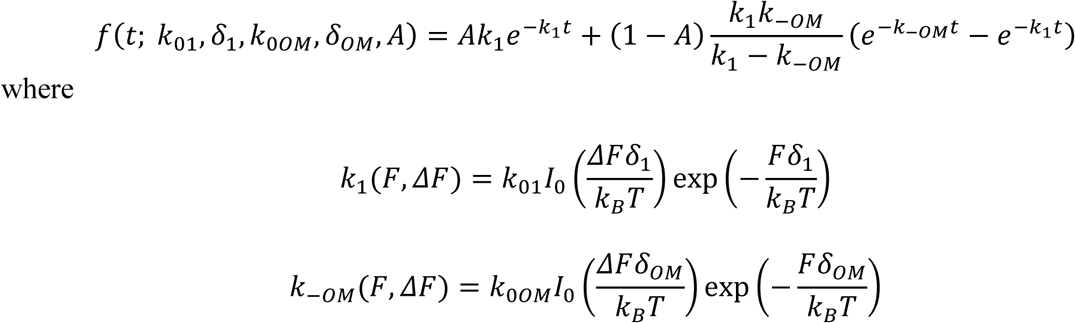

This distribution is a sum of two exponentials, as seen more clearly by redefining the coefficients:

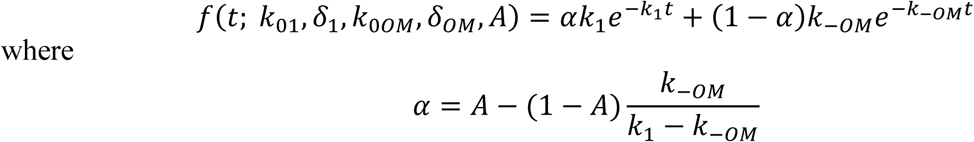

In MLE fitting, the function used is the discrete geometric rather than the continuous exponential distribution due to the requirements of HFS to bin durations in an integer number of periods of stage oscillation^11^. In addition, the minimum detectable time (5 ms) is taken into account by a dead time correction.

MEMLET uses the above probability distribution function to fit to the (*F, ΔF, t_s_*) data, producing the 5 fitted parameters: *k*_01_*, δ*_1_*, k_0OM_, δ_0M_,A.* Here, the advantage of fitting without binning by force can be appreciated: the larger number of fit parameters gives rise to variability when fitting a subset of data in one force bin. For this same reason and for simplicity of analysis, rather than performing separate analyses on individual molecules (as we have done with all other compound and mutation conditions), we aggregated events from all molecules for each OM concentration before performing the MLE. Note that the slow and fast population of events were observed for every single molecule; thus it is not the case that some molecules simply do not bind OM.

At 30 nM OM, the number of events from the slow population is not sufficiently large for an accurate fitting by MEMLET. Therefore, for the 30 nM OM data point only (Fig. 5e-g), we fixed *k*_*o1*_ = 110 s^−1^ and *δ*_*1*_=1.25 nm and performed the MLE to obtain *k*_*0OM*_, δ_oM_, and *A.* Indeed, the fit produced A = 0.96, confirming that there were very few events in the slow population.

### Normalization of in vitro motility results

The range of WT velocities from different protein preparations was 600-1100 nm/s, and this variability has also been observed previously ^50^. Because of this, velocities and *K*_*s*_ of different conditions (compounds and mutations) were normalize-adjusted by each of their WT controls. For example, if condition x had a velocity of vx, and its corresponding WT control had a velocity of vx_control, then the adjusted velocity was vx_adjusted = vx*vWT/vx_control where vwt is the velocity of the WT measured in this study against which everything else is compared.

### Calculations of duty ratio, average force, and average power

Calculations are given by Eqn. 2-5 using values in Table 1. Values and errors referenced from other studies are normalized by each of their corresponding WT protein controls. Errors in *k*_*attach*_are calculated as propagated errors derived from Eqn. 3.

### Code availability

FAST program code use to analyze motility data is available for download at http://spudlab.stanford.edu/fast-for-automatic-motility-measurements/ as described in a previous publication^20^. Custom code used to analyze HFS data is available upon request.

### Data availability

All data supporting the findings of this study are available within the paper and in the extended data set.

## Supplemental text

### Additional details of the HFS method

In HFS, like in other optical trapping methods, the load force applied to myosin upon actin binding is not immediate due to the finite stage velocity and finite time required to stretch the myosin molecule. Here, the value of the average load is reached in one-fourth of the oscillation period T or less, depending on the phase of cycle at which binding occurs. In our experiments, the stage oscillates at 200 Hz, T = 5 ms, so the load is experienced by myosin within 1.25 ms. The power stroke of myosin happens in ~1 ms ^12^, therefore the power stroke of a fraction of events may occur under forces lower or higher than the average force during the entire event. After the power stroke, the myosin molecule in the ADP-bound state continues to experience the oscillatory load. Next, the release of ADP allows a new ATP to bind and release myosin from actin. Since we performed all experiments at saturating (2 mM) ATP conditions, the rate of ATP binding is very fast compared to the rate of ADP release. Because of this, the lifetimes of events in each force bin follows an apparent single exponential distribution parameterized by the ADP release rate. Therefore, the detachment kinetics we measured in no-compound conditions most likely correspond to the load-dependent ADP release rate.

As discussed in Discussion, different perturbations may change the overall detachment kinetics by altering ADP release or other steps in the bound state. Most notably, OM is thought to dramatically delay the power stroke before ADP release. Because we did not measure the details of the bound state (e.g. power stroke), we use the term “detachment rate” to encompass the bound state overall.

## Acknowledgments

The authors thank all members of the Spudich lab, Jongmin Sung, Suman Nag, and Robert McDowell for discussions and edits to the manuscript; Dan Herschlag, Charles Limouse, and Steve Bonilla for discussions on the data analysis and interpretation; Michael Woody for discussions on OM and help with MEMLET; Josh Baker for discussions on attachment vs. detachment rate limited motility velocity; and Fady Malik for discussions on clinically relevant dosage of OM. We thank MyoKardia, Inc. for providing the various small molecule effectors of the human β-cardiac myosin that were derived from their screens, dATP, and bovine actin. This work was funded by NIH grants RO1GM033289 (J.A.S.), RO1HL117138 (J.A.S.), T32GM007276 (C.L.), TL1RR025742 (C.L.), and F32HL124883 (M.K.); Stanford Bio-X fellowship (C.L.); and Stanford School of Medicine Dean’s Postdoctoral Fellowship (D.S.). The content is solely the responsibility of the authors and does not necessarily represent the official view of the National Institutes of Health.

## Author contributions

C.L. performed single molecule experiments and analyzed the data. M.K. performed in vitro motility experiments and analyzed the data. C.L. and D.S. performed ATPase experiments. D.S. analyzed the ATPase data. M.K., D.S., and K.M.R. expressed and purified protein. C.L. and J.A.S. wrote the paper. All authors discussed the data as it evolved and reviewed and edited the paper.

## Author information

J.A.S. is a founder of Cytokinetics, Inc. and MyoKardia, Inc. and a member of their advisory boards. K.M.R. is a member of the MyoKardia scientific advisory board. Correspondence and requests for materials should be addressed to J.A.S.

---

**Extended Data Figure 1:**
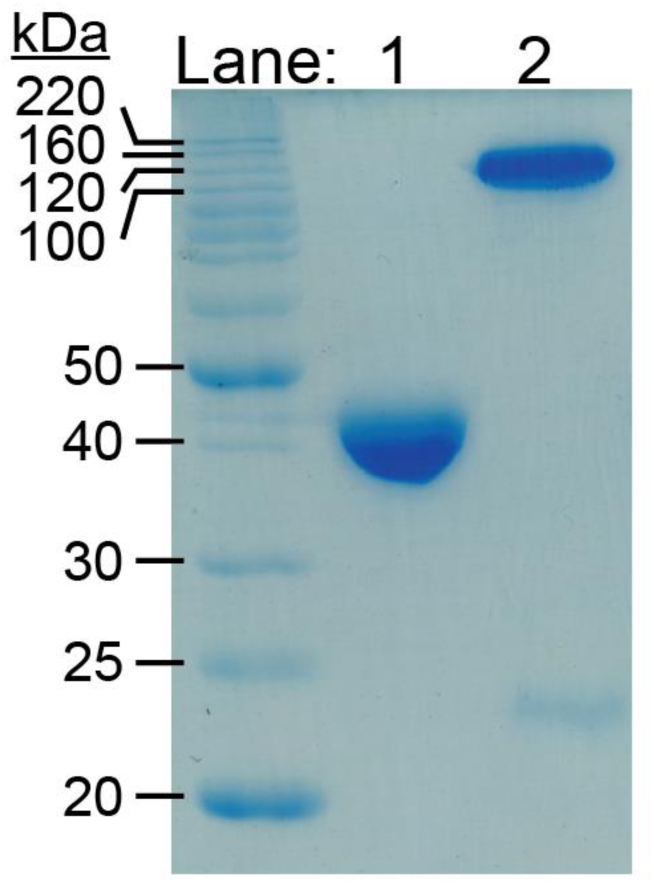
Purified actin and myosin shown by SDS-PAGE. Lane 1 contains bovine cardiac actin (MW = 42 kDa) used in the actin-activated ATPase assay. Lane 2 contains recombinant human β-cardiac myosin sS1 (residue 1-808) fused to a C-terminal eGFP that migrates around 120 kDa. Myosin sS1 fragment was co-purified with a FLAG-tagged human ventricular essential light chain in which the FLAG tag had been cleaved off by TEV protease and migrates around 22 kDa.

**Extended Data Figure 2:**
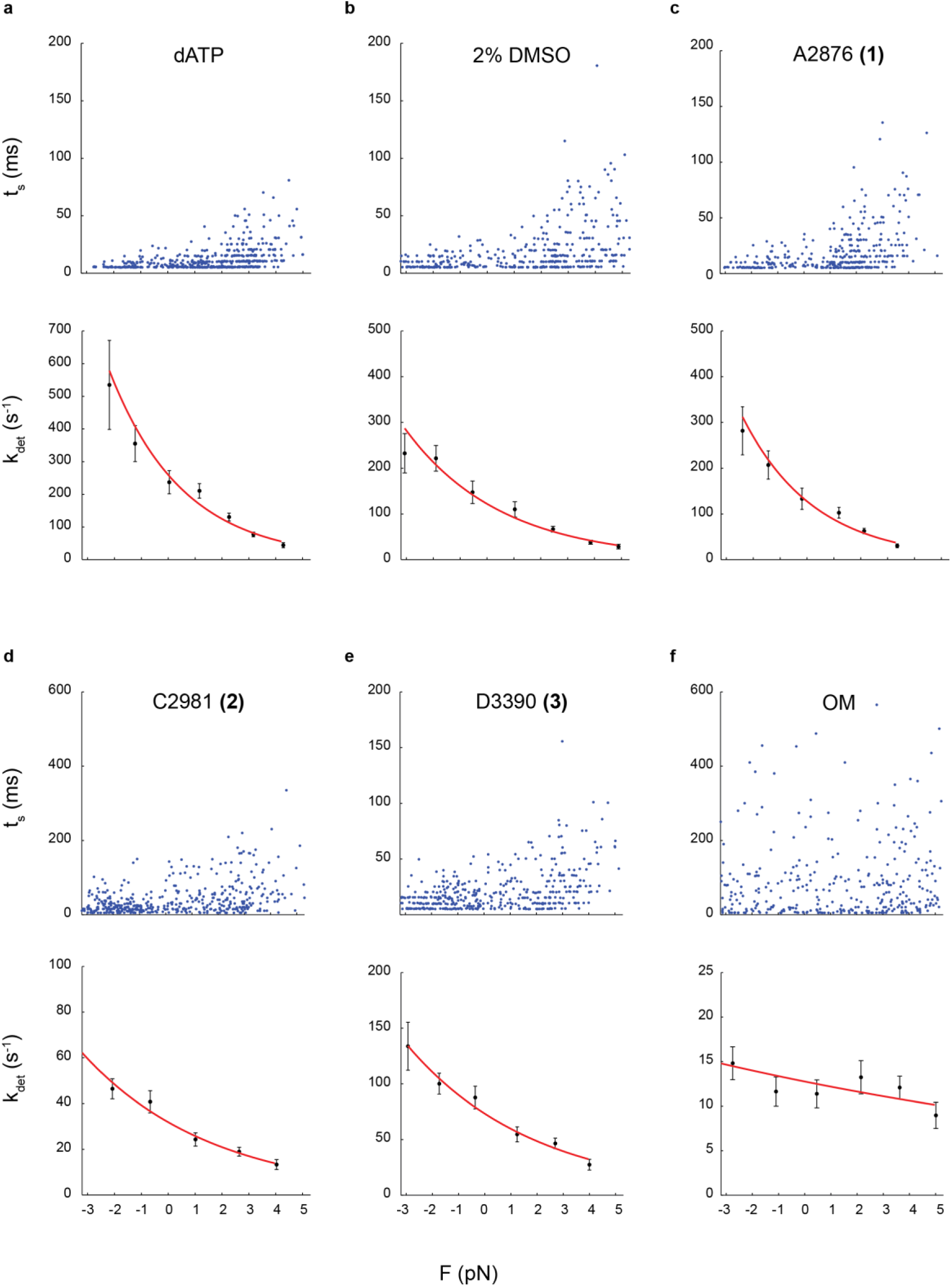
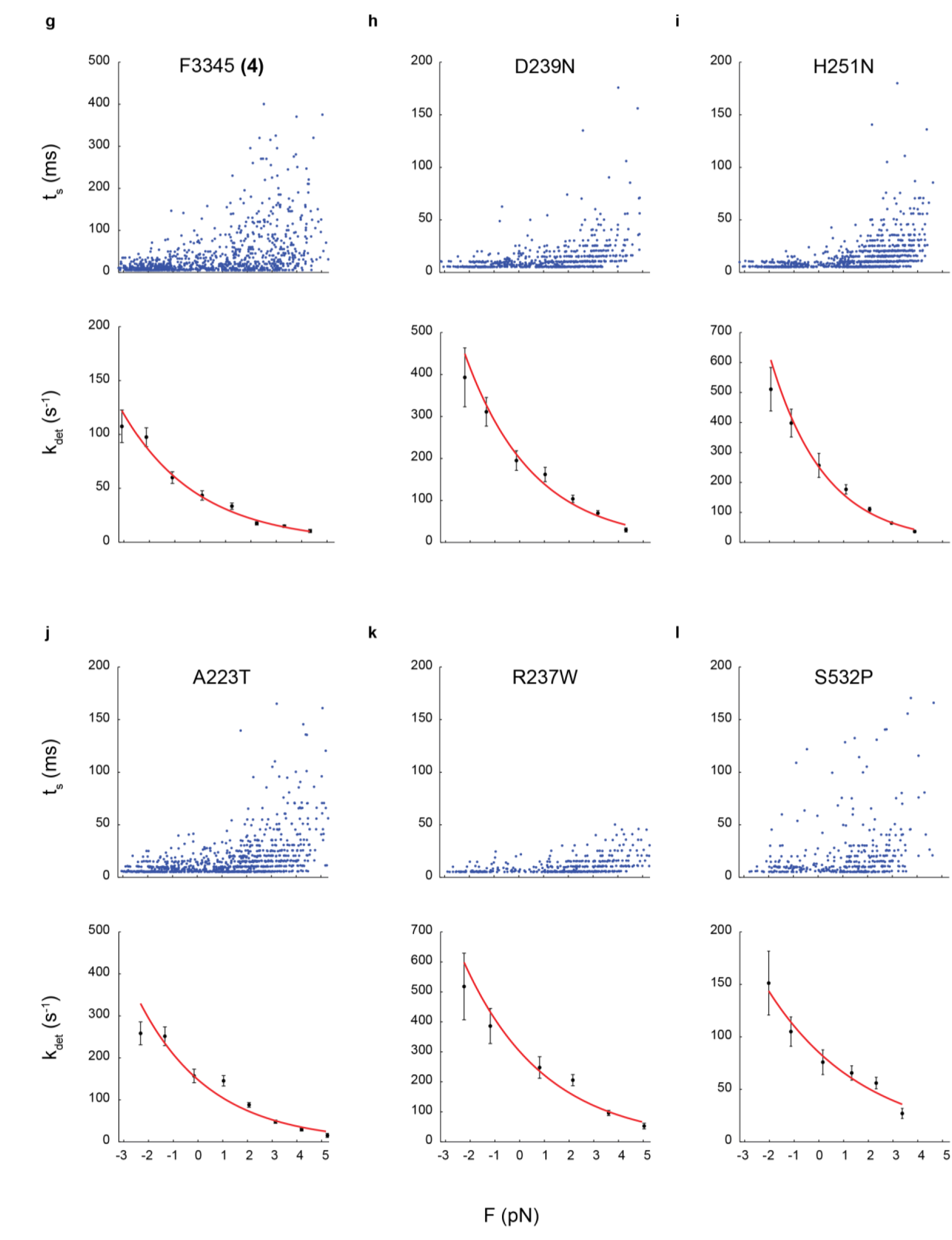
Example molecule from each condition measured. Event lifetimes are plotted against load force (top). The detachment rate *k*_*det*_ at each force is determined by MLE on the exponentially-distributed lifetimes. *k*_*det*_ is then fitted to the Arrhenius equation with harmonic force correction (Eqn. 1) (bottom). An example of WT without compound is shown in Fig. 1c-d.

**Extended Data Figure 3:**
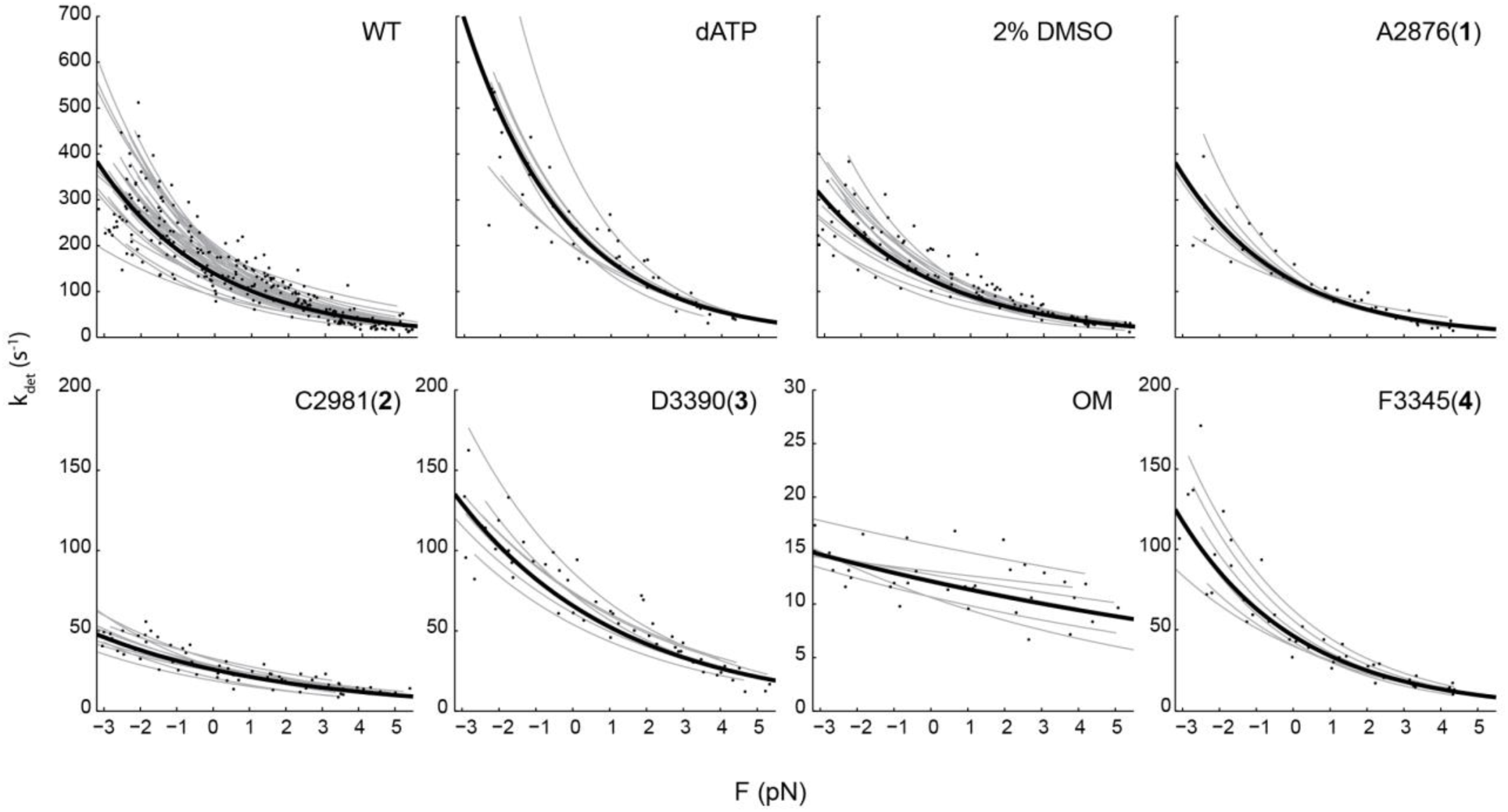
Effects of small molecule compounds on the load-dependent kinetics of single molecules of human β-cardiac myosin. Each gray line represents one molecule fitted to *k*_*det*_ from events binned by force, shown as data points without error bars for clarity. The weighted means of *k*_*0*_ and *δ* across molecules for each condition have a curve represented in black and also plotted in Fig. 3b. Their values are given in Table 1 and Extended Data Table 1. The plot for WT is replicated from Fig. 1f for comparison. In the case of dATP, 2 mM dATP was used in place of ATP. All other conditions used 2 mM ATP, 2% DMSO, and 25 μM compound concentration.

**Extended Data Figure 4:**
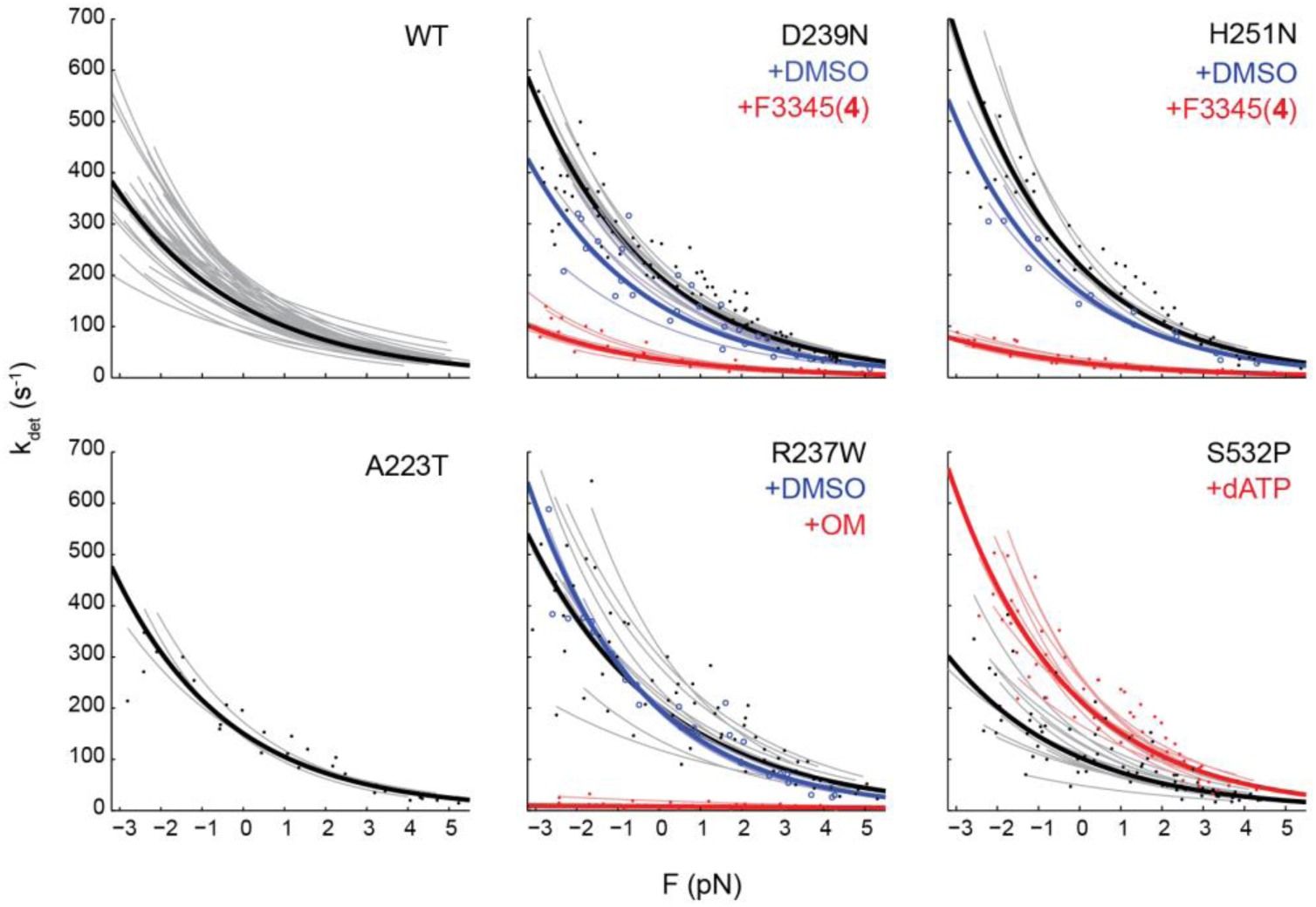
Effects of cardiomyopathy-causing mutations on the load-dependent kinetics of single molecules of human β-cardiac myosin, and their reversal by small molecule compounds. Each thin line represents one molecule fitted to *k*_*det*_ from events binned by force, shown as data points without error bars for clarity. The weighted means of *k*_*0*_ and *δ* across molecules for each condition have a curve represented as a thick line and also plotted in Fig. 4b. Their values are given in Table 1 and Extended Data Table 1. Addition of 25 μM compound F3345 (4) to D239N and H251N and OM to R237W were in the presence of 2% DMSO, therefore effects of 2% DMSO (blue) on these mutants were also measured. Mutant A223T had no significant change from WT, therefore no compounds were added. The plot for WT is replicated from Fig. 1f for comparison.

**Extended Data Figure 5:**
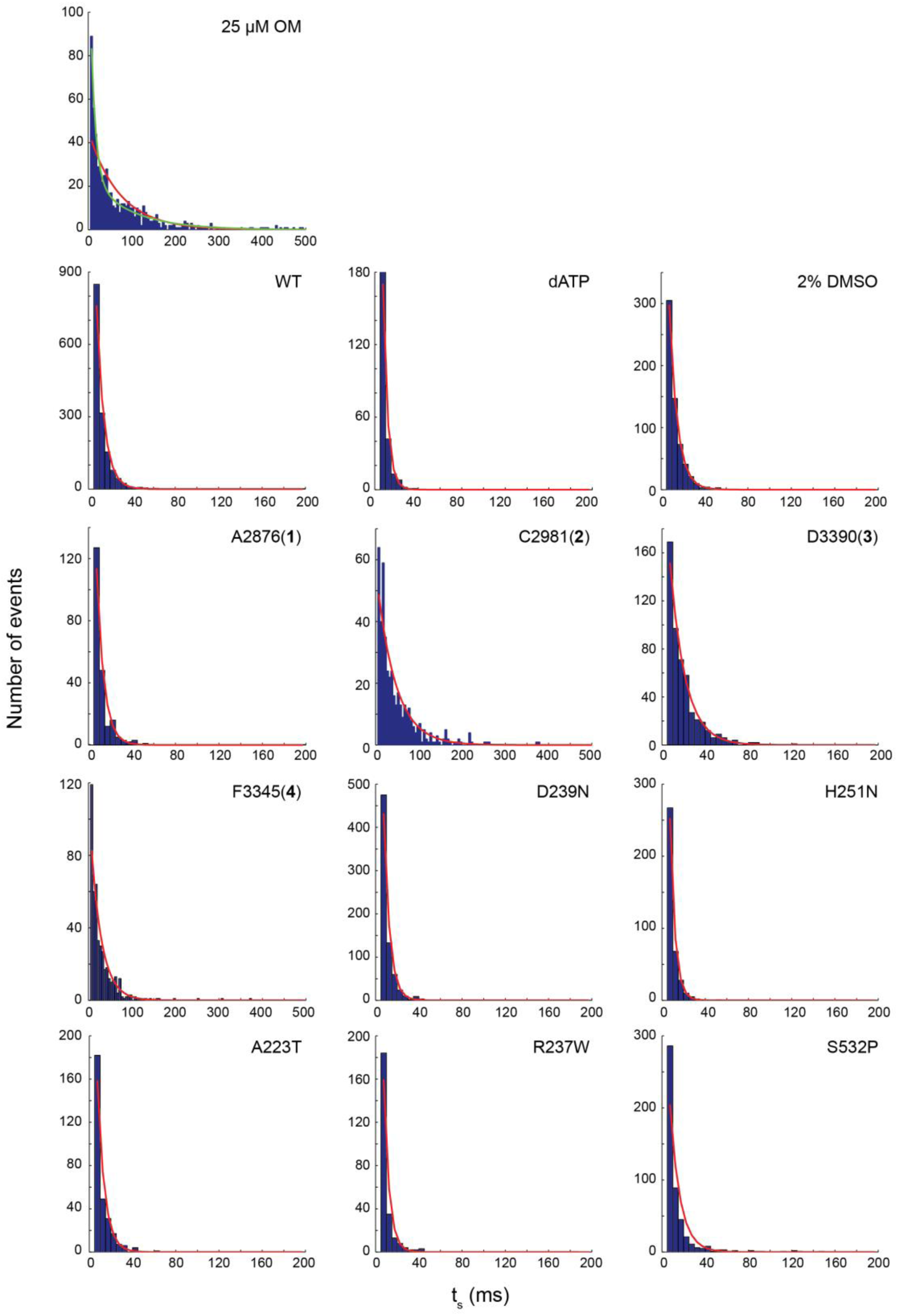
Durations of all events in the F = 0 force bin from all molecules for each condition measured. For each condition, the red curve represents the single exponential distribution parameterized by a single detachment rate as determined by MLE. Even at the saturating 25 μM concentration of OM, there are still two populations better described by a double (green) rather than a single (red) exponential distribution (top). F ≠ 0 bins show similar single (all conditions except OM) or double (OM) exponential distributions (not shown).

**Extended Data Figure 6:**
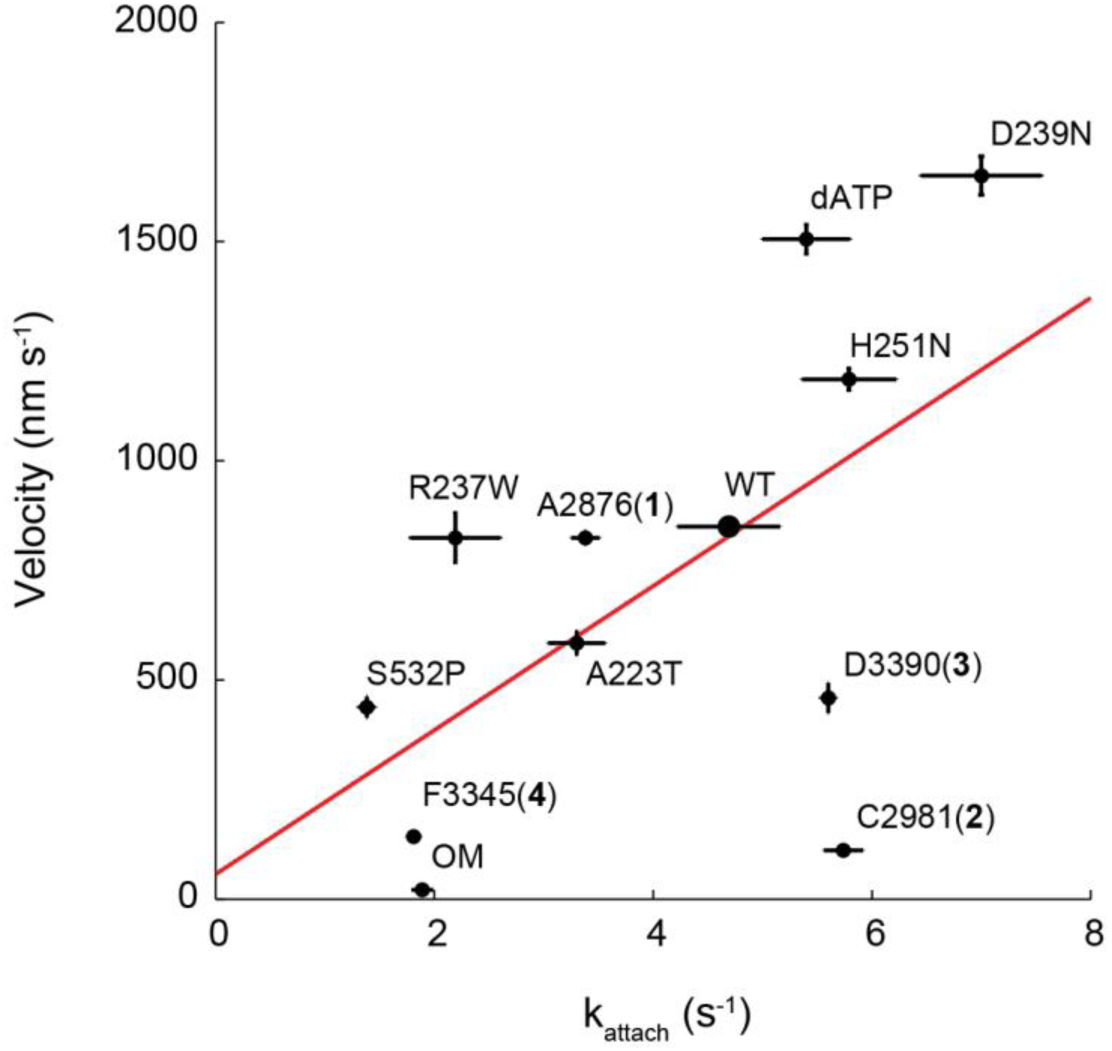
Unloaded actin sliding velocity measured by in vitro motility assay vs. the attachment rate *k*_*attach*_ calculated by Eqn. 3. Linear regression gives R^2^ = 0.35. Motility velocity of HCM, DCM, and dATP are obtained from other studies^20,25,32^ (Ujfalusi Z., Vera C., Mijailovich S., Svicevic M., Choe Yu E., Kawana M., Ruppel K., Spudich J., Geeves M., and Leinwand L., *JBC* manuscript under review; Tomasic I., Liu C., Rodriguez H., Spudich J.A., Bartholomew Ingle S.R., manuscript in preparation).

**Extended Data Figure 7:**
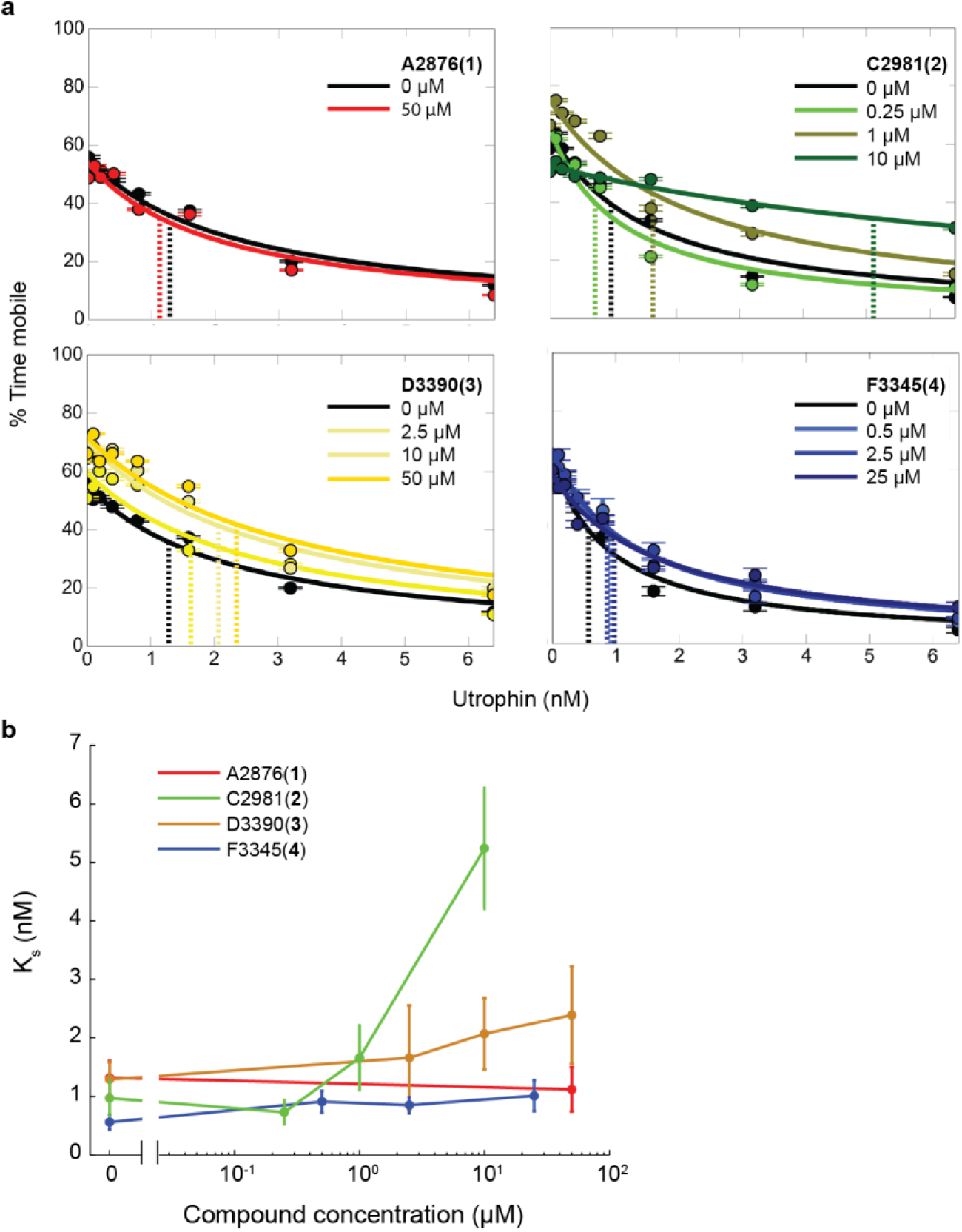
Effects of compounds on the actin-sliding velocities of human β-cardiac myosin in the loaded in vitro motility. **a**, Percent time mobile (a measure of velocity) as a function of the actin-binding protein utrophin, which served as a resistive load against myosin in the motility assay, as different concentrations of compound are added to WT myosin. Error bars represent s.e.m. of bootstrapped data calculated by the FAST program ^20^. Dashed lines denote *K_s_,* a measure of the ensemble load-bearing ability of myosin. *K*_*s*_ roughly corresponds to the concentration of utrophin required to slow down velocity by half (see ^20^ for rigorous definition). **b**, *K*_*s*_ from (a) is plotted against compound concentration. Error bars represent s.e.m.

**Extended Data Figure 8:**
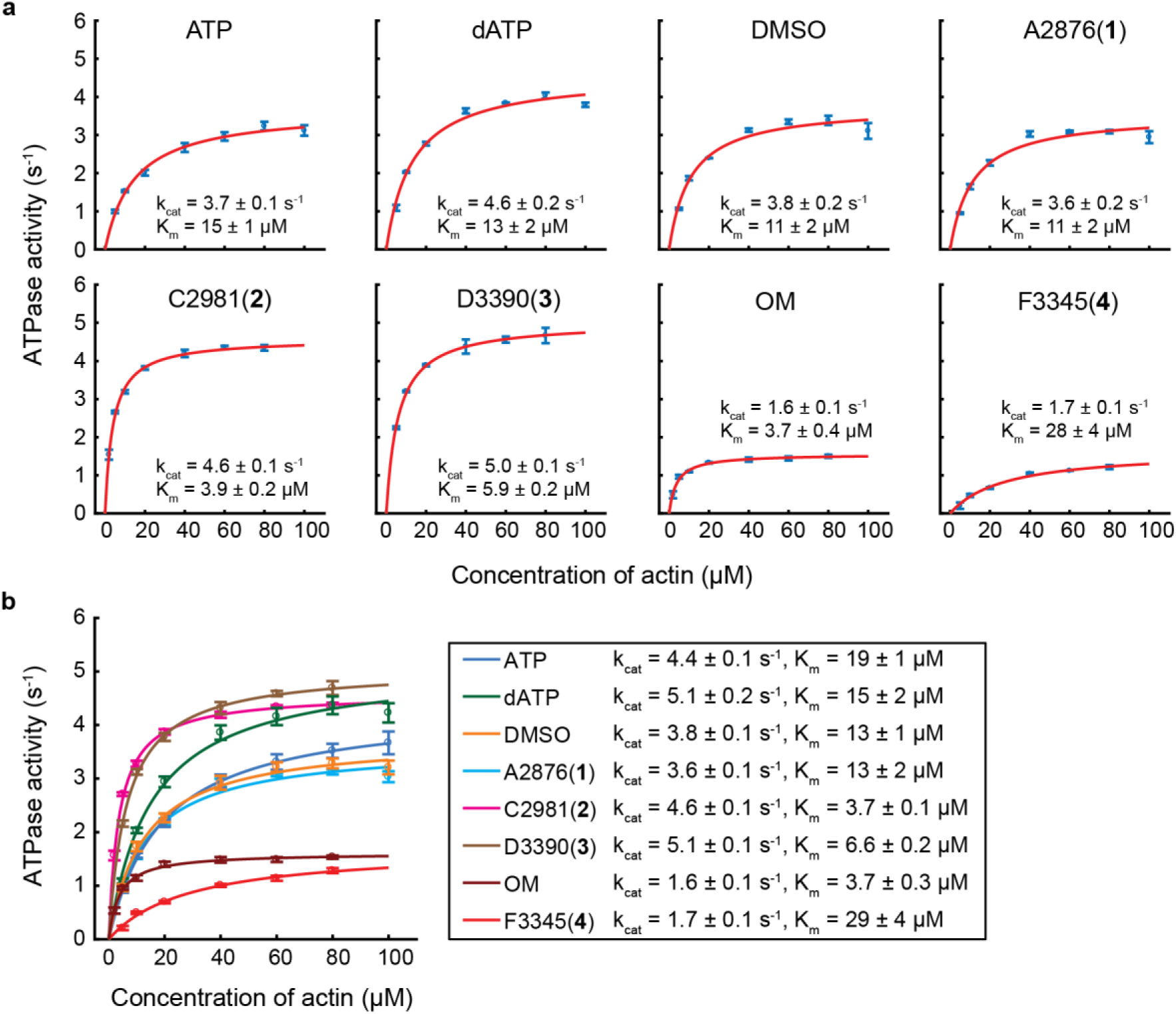
Effects of compounds on the actin-activated ATPase of human β-cardiac myosin. **a**, The Michaelis-Menten equation fitted to ATPase data for all conditions performed with one protein preparation. The entire experiment measuring all conditions was done on the same day that the protein was purified. Error bars on data points are s.e.m. of replicate experiments (N=3 in this example). Errors on *k*_*cat*_ and *K*_*m*_ are estimated fitting errors. *k*_*eat’s*_ from each day’s experiment are averaged to calculate the mean and s.e.m. given in Table 1. **b**, The Michaelis-Menten equation fitted to ATPase data aggregated from both protein preparations. Here, error bars on data points are s.e.m. of all replicates (total 5 - 7), and errors on *k*_*cat*_ and *K*_*m*_ are estimated fitting errors.

**Extended Data Figure 9:**
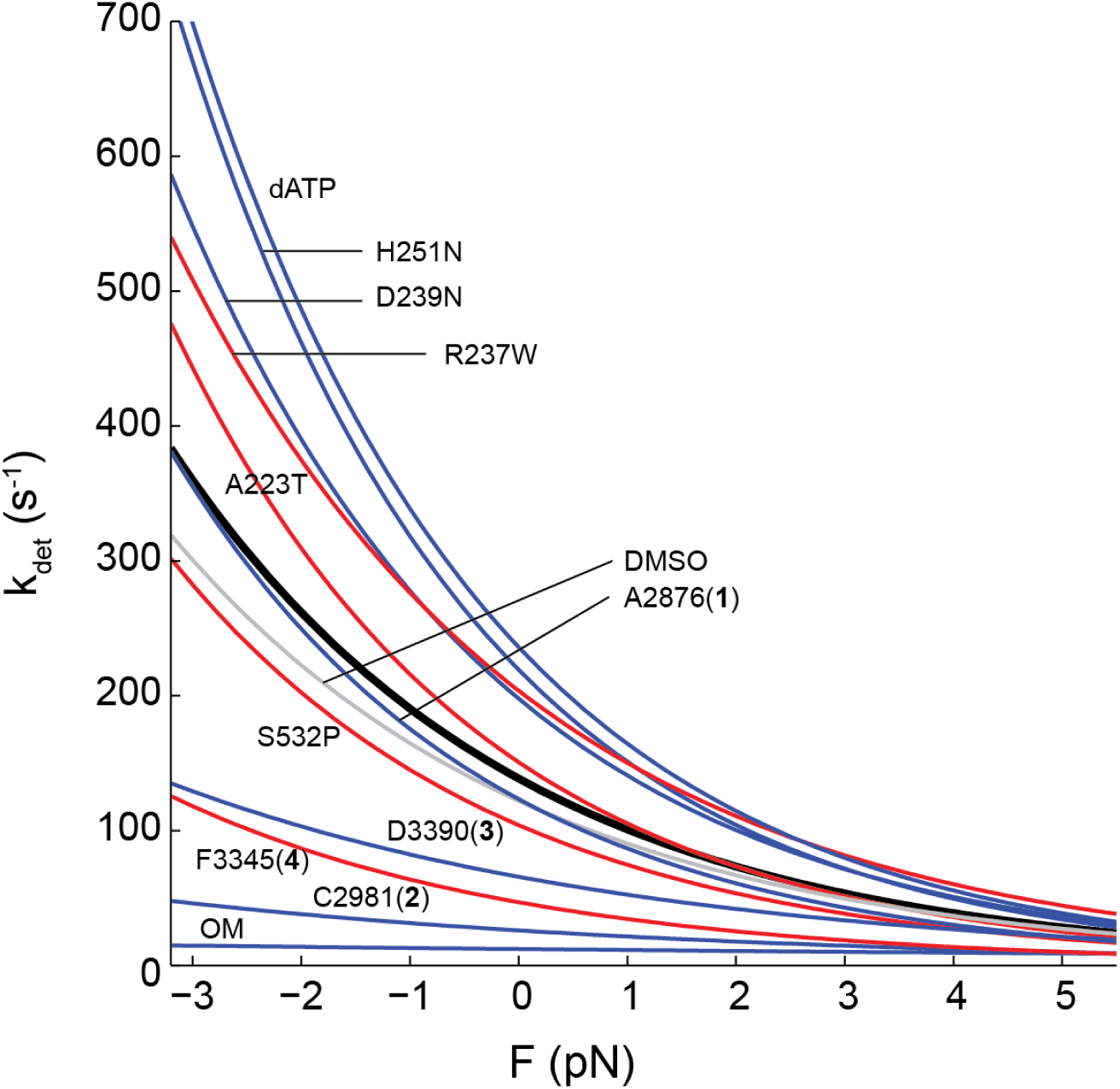
Small molecule compounds and cardiomyopathy-causing mutations modulate the load-dependent kinetics of single cardiac myosin molecules across a spectrum. The weighted average detachment rate curves for the different conditions are replicated from Figs. 3 and 4. Their *k*_*0*_ and *δ* values are given in Table 1. Black: WT. Gray: DMSO. Blue: HCM and activators. Red: DCM and inhibitor.

**Extended Data Table 1:** Summary of single molecule detachment kinetics data (*k*_*0*_,*δ*) for human β-cardiac myosin with effects of small molecule compounds and cardiomyopathy-causing mutations. Values are mean ± s.e.m. In the case of dATP, 2 mM dATP was used in place of ATP. All other conditions used 2 mM ATP, 2% DMSO, and 25 µM compound concentration. 2-tailed unequal variances t-test performed on dATP vs WT (ATP), DMSO vs WT, compounds + DMSO vs DMSO alone, mutants vs WT, and mutants + compounds + DMSO vs mutants + DMSO alone.

| | $k_0$ (s<sup>-1</sup>) | p value | $\delta$ (nm) | p value | # molecules |
| --- | --- | --- | --- | --- | --- |
| **WT** | 102 ± 4 |  | 1.31 ± 0.03 |  | 36 |
| **+dATP** | 168 ± 7 | 3.5e-5 | 1.49 ± 0.10 | 0.13 | 7 |
| **+DMSO** | 87 ± 4 | 1.4e-4 | 1.24 ± 0.04 | 0.59 | 16 |
| **+A2876 (1)** | 87 ± 2 | 0.60 | 1.45 ± 0.13 | 0.18 | 6 |
| **+C2981 (2)** | 23 ± 1 | 1.1e-12 | 0.79 ± 0.05 | 1.0e-7 | 10 |
| **+D3390 (3)** | 55 ± 3 | 1.7e-6 | 0.93 ± 0.03 | 7.7e-7 | 7 |
| **+OM** | 12 ± 1 | 4.0e-13 | 0.26 ± 0.05 | 4.0e-10 | 6 |
| **+F3345 (4)** | 36 ± 2 | 1.2e-10 | 1.27 ± 0.06 | 0.8 | 6 |
| **D239N** | 140 ± 3 | 3.1e-7 | 1.40 ± 0.03 | 0.04 | 12 |
| **+DMSO** | 108 ± 14 | 0.19 | 1.41 ± 0.04 | 0.74 | 5 |
| **+F3345 (4)** | 24 ± 2 | 0.0026 | 1.35 ± 0.08 | 0.71 | 6 |
| **H251N** | 152 ± 7 | 4.8e-4 | 1.53 ± 0.08 | 0.02 | 6 |
| **+DMSO** | 119 ± 4 | 0.0039 | 1.51 ± 0.09 | 0.60 | 2 |
| **+F3345 (4)** | 20 ± 2 | 0.0050 | 1.27 ± 0.05 | 0.17 | 5 |
| **A223T** | 104 ± 4 | 0.17 | 1.48 ± 0.09 | 0.13 | 4 |
| **R237W** | 155 ± 15 | 0.0039 | 1.25 ± 0.06 | 0.43 | 10 |
| **+DMSO** | 142 ± 4 | 0.12 | 1.51 ± 0.07 | 0.018 | 4 |
| **+OM** | 8 ± 1 | 2.4e-5 | 0.21 ± 0.06 | 1.0e-6 | 9 |
| **S532P** | 80 ± 8 | 0.021 | 1.37 ± 0.06 | 0.57 | 12 |
| **+dATP** | 170 ± 6 | 1.4e-7 | 1.45 ± 0.08 | 0.12 | 9 |

